# pH-responsive upconversion mesoporous silica nanospheres for combined multimodal diagnostic imaging and targeted photodynamic and photothermal cancer therapy

**DOI:** 10.1101/2023.05.22.541491

**Authors:** L. Palanikumar, Mona Kalmouni, Tatiana Houhou, Osama Abdullah, Liaqat Ali, Renu Pasricha, Sneha Thomas, Ahmed J. Afzal, Francisco N. Barrera, Mazin Magzoub

**Affiliations:** Biology Program, Division of Science, New York University Abu Dhabi, Abu Dhabi, United Arab Emirates; Core Technology Platforms, New York University Abu Dhabi, Abu Dhabi, United Arab Emirates; Department of Biochemistry & Cellular and Molecular Biology, University of Tennessee Knoxville, Knoxville, Tennessee, United States

**Keywords:** ATRAM peptide, cancer therapy, mesoporous silica, magnetic resonance imaging, near-infrared light, photodynamic therapy, photothermal therapy, reactive oxygen species, tumor microenvironment, upconversion

## Abstract

Photodynamic therapy (PDT) and photothermal therapy (PTT) have garnered considerable interest as non-invasive cancer treatment modalities. However, these approaches remain limited by low solubility, poor stability and inefficient targeting of many common photosensitizers (PSs) and photothermal agents (PTAs). To overcome these limitations, we have designed biocompatible and biodegradable tumor-targeted upconversion nanospheres with imaging capabilities. The multifunctional nanospheres consist of a sodium yttrium fluoride core doped with lanthanides (ytterbium, erbium and gadolinium) and bismuth selenide (NaYF_4_:Yb/Er/Gd,Bi_2_Se_3_) within a mesoporous silica shell that encapsulates a PS, Chlorin e6 (Ce6), in its pores. NaYF_4_:Yb/Er converts deeply penetrating near-infrared (NIR) light to visible light, which excites the Ce6 to generate cytotoxic reactive oxygen species (ROS), while the PTA Bi_2_Se_3_ efficiently converts absorbed NIR light to heat. Additionally, Gd enables magnetic resonance imaging (MRI) of the nanospheres. The mesoporous silica shell is coated with lipid/polyethylene glycol (DPPC/cholesterol/DSPE-PEG) to ensure retention of the encapsulated Ce6 and minimize interactions with serum proteins and macrophages that impede tumor targeting. Finally, the coat is functionalized with the acidity-triggered rational membrane (ATRAM) peptide, which promotes specific and efficient internalization into cancer cells within the mildly acidic tumor microenvironment. Following uptake by cancer cells *in vitro*, NIR laser irradiation of the nanospheres caused substantial cytotoxicity due to ROS production and hyperthermia. The nanospheres facilitated tumor MRI and thermal imaging, and exhibited potent NIR laser light-induced antitumor effects *in vivo* via combined PDT and PTT, with no observable toxicity to healthy tissue, thereby substantially prolonging survival. Our results demonstrate that the ATRAM-functionalized, lipid/PEG-coated upconversion mesoporous silica nanospheres (ALUMSNs) offer multimodal diagnostic imaging and targeted combinatorial cancer therapy.

## INTRODUCTION

Traditional cancer treatments, such as chemotherapy, radiotherapy and surgery, suffer from a number of issues that severely limit their clinical efficacy, including a range of side-effects and complications, immunosuppression, development of multidrug resistance (MDR) phenotypes, recurrence and metastasis^1–3^. This has created a pressing need for new therapeutic strategies to supplement or supplant conventional cancer treatments. Foremost among these alternatives are non-invasive light-based therapies, photodynamic therapy (PDT) and photothermal therapy (PTT), which have gained considerable attention as potentially safe and effective modalities^4, 5^. PDT uses laser irradiation to activate a photosensitizer (PS) that subsequently generates cytotoxic reactive oxygen species (ROS), through a series of photochemical reactions, to induce apoptosis in cancer cells, while in PTT a photothermal agent (PTA) converts absorbed light into heat and the resulting hyperthermia leads to the partial or complete ablation of tumor tissue^5, 6^.

Despite its therapeutic potential, PDT currently has several drawbacks. The majority of PS molecules are cyclic tetrapyrroles, and many of these are characterized by poor solubility, rapid *in vivo* degradation and clearance, and lack of tumor specificity^5^. These characteristics are particularly problematic given that ROS is highly reactive and consequently has a very short lifetime (< 40 ns) and limited radius of action (∼20 nm) in cellular milieu, which necessitates accumulation of PSs within tumors for effective PDT^7^. Furthermore, since PSs use molecular oxygen to generate ROS, the hypoxic microenvironment of tumors can greatly impair PDT^8, 9^. Likewise, PTT has a number of issues. PTAs are divided into inorganic (e.g., gold nanoparticles, transition metal sulfides or oxides, graphene and carbon nanotubes) and organic (e.g., cyanine, porphyrin, diketopyrrolopyrrole and polymers) materials^10–12^. Inorganic PTAs are generally characterized by poor biocompatibility and biodegradability, whereas most organic PTAs exhibit low photothermal conversion efficiency and photostability and neither group possesses inherent tumor specificity^5, 11, 13^. Another challenge is that hyperthermia often leads to overexpression of heat shock proteins, as part of the stress response, which confers a degree of thermotolerance to cancer cells that diminishes the effects of PTT^14, 15^. Temperatures > 50 °C are therefore necessary to overcome this acquired thermotolerance and achieve complete tumor ablation (via protein denaturation and plasma membrane destruction), but the high irradiation power densities and longer illumination durations necessary to reach these elevated temperatures also pose a risk of irreversible damage to the surrounding non-malignant tissue due to the presence of endogenous chromophores^15^.

The current limitations of PDT and PTT have meant that neither form of phototherapy alone is sufficient to completely eradicate tumors^16^. This has prompted the development of nanocarriers for more effective tumor delivery of PS and PTA molecules^5, 16^. A particularly promising strategy is nanocarrier-mediated simultaneous delivery of PSs and PTAs as means of combining the two forms of phototherapy in order to synergistically enhance their antitumor effect^5^. The advantage of this approach is that PTT-induced hyperthermia can facilitate accumulation of PS molecules and molecular oxygen in tumor tissue by boosting local blood flow, which serves to improve the effectiveness of PDT, while ROS produced during PDT can inhibit heat shock proteins, thereby decreasing the thermotolerance of cancer cells and increasing their susceptibility to PTT^17–19^. However, the complex modifications often used to incorporate PS and PTA molecules into the same nanocarrier can attenuate the therapeutic efficacy of the system^20^. Furthermore, currently < 1% of intravenously administered NPs accumulate in solid tumors^21, 22^. This can be attributed, in large part, to the formation of a serum protein corona on the surface of nanocarriers during *in vivo* circulation^23^. The adsorbed serum proteins not only destabilize nanocarriers, but also trigger immune recognition and rapid blood clearance, all of which culminates in poor tumor accumulation^24^. Finally, for the small fraction of nanocarriers that does reach the target tumor tissue, uptake into cancer cells represents a major challenge. The primary cellular internalization route for the majority of nanocarriers is endocytosis, but endosomal escape efficiency remains extremely low (1–2%), with most endocytosed nanocarriers becoming entrapped in degradative acidic endocytic compartments or undergoing exocytosis^22, 25^.

Here, we have developed multifunctional nanospheres that overcome the aforementioned challenges. These biocompatible and biodegradable core-shell nanospheres consist of a lanthanide- and PTA-doped upconversion core within a PS-loaded mesoporous silica shell. The shell is wrapped with a lipid/PEG bilayer that is functionalized with the tumor targeting acidity-triggered rational membrane (ATRAM) peptide^26^. The ATRAM-functionalized, lipid/PEG-coated upconversion mesoporous silica nanospheres (ALUMSNs) enable tumor detection via MRI and thermal imaging. The ALUMSNs additionally facilitate NIR laser light-induced PDT and PTT to substantially shrink tumors, with no detectable adverse effects to healthy tissue, leading to markedly prolonged survival.

## RESULTS AND DISCUSSION

### Preparation and characterization of upconversion mesoporous silica nanospheres (UMSNs)

The core of the upconversion mesoporous silica nanospheres (UMSNs) consists of sodium yttrium fluoride doped with lanthanides (ytterbium, erbium and gadolinium) and bismuth selenide (NaYF_4_:Yb/Er/Gd,Bi_2_Se_3_) (Figure 1a). Transmission electron microscopy (TEM) and scanning transmission electron microscopy (STEM) images showed a uniform sphere-like upconversion core (Figure 2a; Supporting Figure 1a, b). The composition of the upconversion core was verified using scanning transmission electron microscopy-energy dispersive X-ray spectroscopy (STEM-EDS) mapping (Supporting Figure 1c). The average hydrodynamic diameter of the core was confirmed by dynamic light scattering (DLS) to be ∼60 nm (Figure 2d).

**Figure 1.**
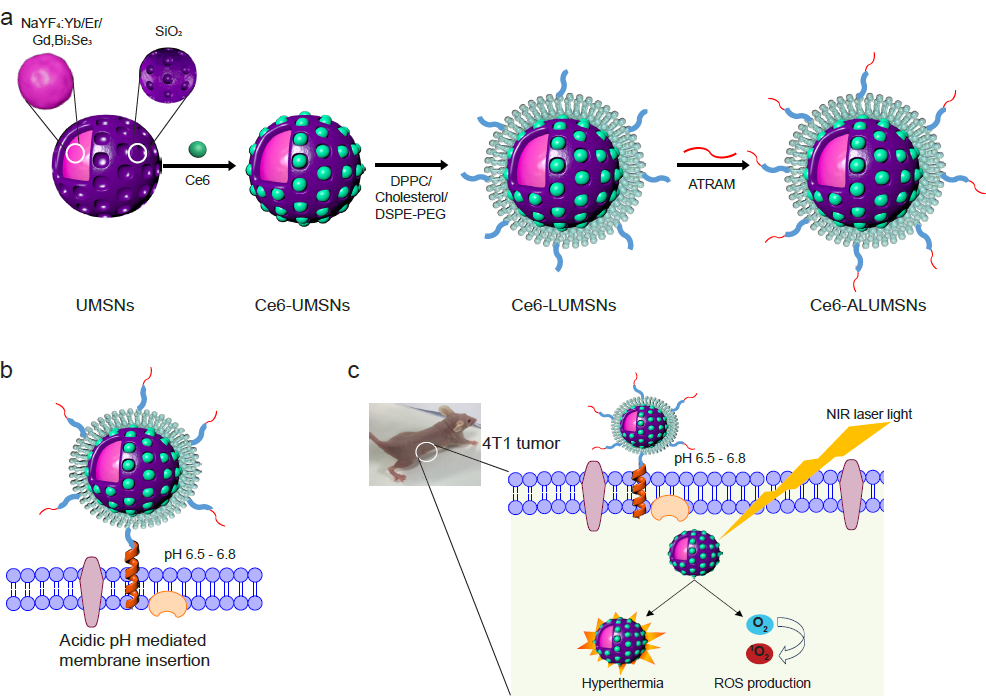
Schematic representation of preparation and mode of action of tumor-targeted upconversion mesoporous silica nanospheres. (**a**) The nanospheres consist of an upconversion core of sodium yttrium fluoride doped with lanthanides – ytterbium, erbium, and gadolinium – and bismuth selenide (NaYF_4_:Yb/Er/Gd,Bi_2_Se_3_) within a mesoporous silica shell that encapsulates a photosensitizer, Chlorin e6 (Ce6), in its pores. The Ce6-loaded upconversion mesoporous silica nanospheres (Ce6-UMSNs) are then ‘wrapped’ with lipid/polyethylene glycol (DPPC/cholesterol/ DSPE-PEG_2000_-maleimide). Finally, the Ce6-loaded lipid/PEG coated UMSNs (Ce6-LUMSNs) are functionalized with the acidity-triggered rational membrane (ATRAM) peptide. (**b**) In mildly acidic conditions, ATRAM inserts into lipid bilayers as a transmembrane α-helix. As the membrane insertion p*K*_a_ of ATRAM is 6.5^26^, the peptide promotes targeting of ATRAM-functionalized Ce6-LUMSNs (Ce6-ALUMSNs) to cancer cells in the mildly acidic (pH ∼6.5–6.8) microenvironment of solid tumors^90^. (**c**) ALUMSNs are efficiently internalized into tumor cells, where subsequent NIR (980 nm) laser irradiation of the nanospheres results in substantial cytotoxicity due to the combined effects of hyperthermia and reactive oxygen species (ROS) generation.

**Figure 2.**
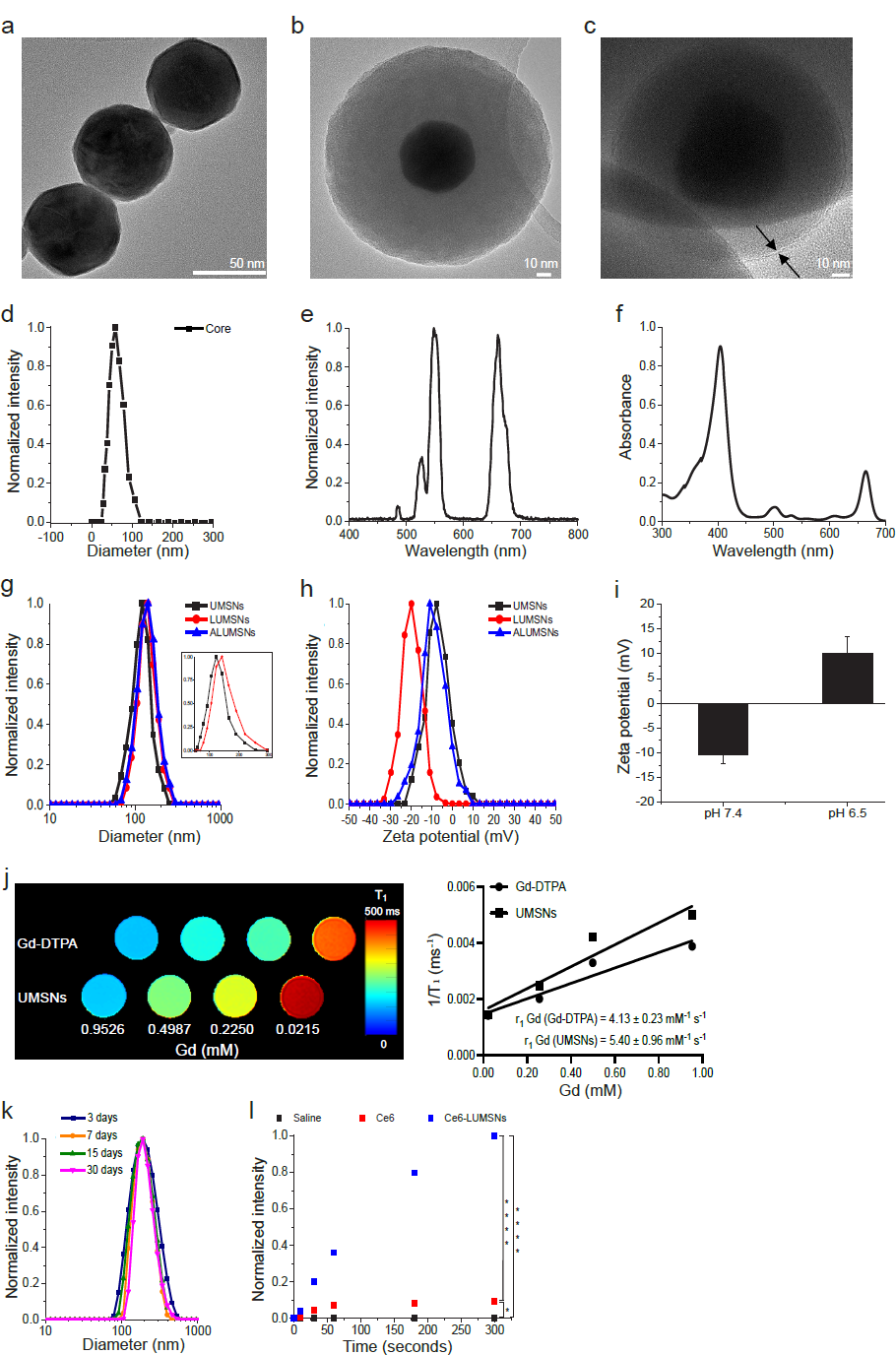
Characterization of the upconversion mesoporous silica nanospheres. (**a**–**c**) Transmission electron microscopy (TEM) images of the upconversion core (NaYF_4_:Yb/Er/Gd,Bi_2_Se_3_) (**a**), upconversion mesoporous silica nanoparticles (UMSNs) (**b**) and lipid/PEG-coated UMSNs (LUMSNs) (**c**). The arrows in (**c**) indicate the lipid bilayer. Scale bar in (**a**) = 50 nm, in (**b**) and (**c**) = 10 nm. (**d**) Size analysis for the upconversion core in 10 mM phosphate buffer (pH 7.4) using dynamic light scattering (DLS). (**e**) Fluorescence emission spectrum of the upconversion core (*λ*_ex_ = 980 nm). (**f**) UV-Vis absorption spectrum of Chlorin e6 (Ce6) (Soret peak at 404 nm and Q-band at 658 nm)^33^. (**g**,**h**) Size distribution analysis (**g**) and zeta potential measurements (**h**) for UMSNs, LUMSNs and ALUMSNs in 10 mM phosphate buffer (pH 7.4). *Inset:* expanded scale to show difference in hydrodynamic diameters of UMSNs and LUMSNs. (**i**) Comparison of zeta potentials of ATRAM-functionalized LUMSNs (ALUMSNs) at pHs 7.4 and 6.5. (**j**) T_1_ maps (*left*) and the relaxation rates (1/T_1_) (*right*) of UMSNs compared to commercial Gd-DTPA (at the same concentrations of the lanthanide) (*n* = 3). (**k**) Colloidal stability analysis for LUMSNs in complete cell culture medium (RPMI 1640 containing 10% fetal bovine serum (FBS), pH 7.4) over 30 days at 37 °C. (**l**) Comparison of ROS production capability of Ce6-LUMSNs and free Ce6, at the same Ce6 concentration (0.5 µg/mL) and NIR laser irradiation power density and duration (980 nm, 1.0 W/cm^2^, 5 min), monitored in 10 mM phosphate buffer (pH 7.4) using the fluorescent probe Singlet Oxygen Sensor Green (SOSG)^52, 53^. * *P* < 0.05, **** *P* < 0.001 for comparisons with controls or amongst the different samples.

Using a facile synthesis method^27^, the core was enveloped in a mesoporous silica shell (Figure 1a). Mesoporous silica was selected due to its physicochemical properties that make it highly suited for drug delivery applications: excellent biocompatibility and biodegradability, high thermal and chemical stability, large surface area for drug loading by adsorption, tunable pore size to modulate drug release kinetics, and ease of surface modification for increased *in vivo* circulation time and improved targeting^28, 29^. N_2_ adsorption-desorption isotherms showed that the shell has a specific surface area of ∼700 m^2^/g and an average pore diameter of ∼2.3 nm (Supporting Figure 2a), which is within the range reported for other promising mesoporous silica-based drug delivery nanoplatforms^30, 31^. TEM, high-angle annular dark-field scanning transmission electron microscopy (HAAD-STEM), and STEM-EDS confirmed the formation of UMSNs as a core-shell structure, which had a hydrodynamic diameter of 160 ± 10 nm and a zeta potential of –6 mV (Figure 2b, g, h; Supporting Figure 2b–d; Supporting Table 1). The photosensitizer (PS) Chlorin e6 (Ce6) was encapsulated within the pores of the mesoporous silica shell using a passive entrapment loading technique. By adjusting the feed ratio, a relatively high loading capacity of Ce6 in the UMSNs was achieved (22 wt%; Supporting Table 2).

The core NaYF_4_:Yb/Er is excited by near-infrared (NIR) light, which has greater tissue penetration, lower autofluorescence and reduced phototoxicity compared to visible light^32^. Spectroscopic analysis revealed clear overlap between the fluorescence emission of the upconversion core and the absorption of Ce6 at the Q-band at 658 nm (Figure 2e,f)^33^. Therefore, under NIR irradiation, the fluorescence emission from the upconverting core will excite the Ce6 encapsulated within the pores of the UMSNs to generate cytotoxic reactive oxygen species (ROS) (Supporting Figure 3). The photothermal agent (PTA) Bi_2_Se_3_ was additionally incorporated in the core to simultaneously convert the absorbed NIR light to heat for photothermal therapy and imaging^34^. Finally, by doping the core with Gd, UMSNs can also serve as MRI contrast agents. T_1_ maps and relaxation rate (1/T_1_) plots^35, 36^ demonstrate that UMSNs consistently yielded greater contrast enhancement compared to the clinically used contrast agent Gd-DTPA (at the same concentrations of the lanthanide; Figure 2j).

**Figure 3.**
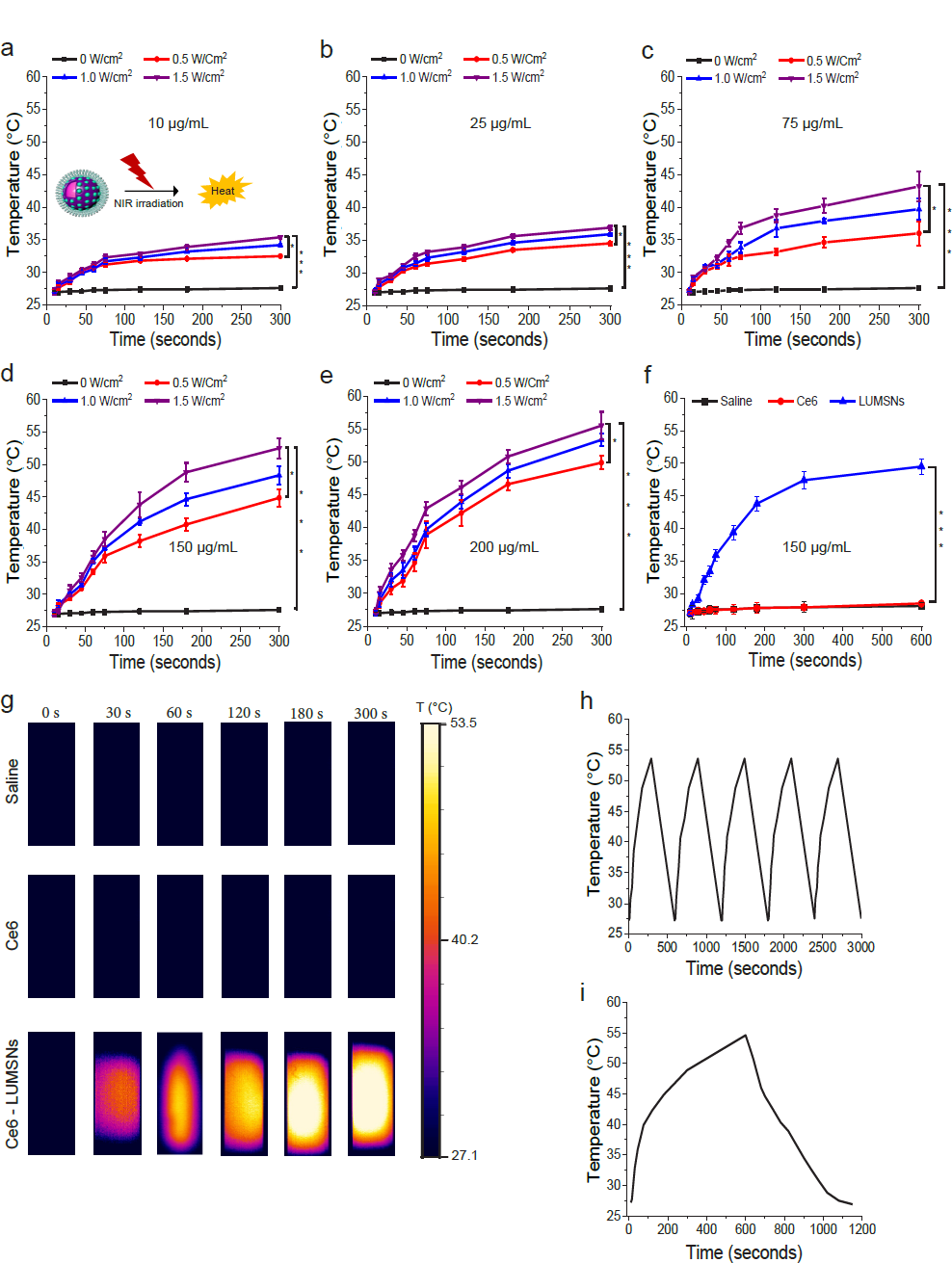
Photothermal properties of Ce6-loaded LUMSNs. (**a**–**e**) Temperature increases following NIR laser irradiation (0.5–1.5 W/cm^2^, 5 min) of Ce6-LUMSNs at nanosphere concentrations of 10 µg/mL (**a**), 25 µg/mL (**b**), 75 µg/mL (**c**), 150 µg/mL (**d**) and 200 µg/mL (**e**) Ce6-LUMSNs in 10 mM phosphate buffer (pH 7.4). (**f**) Comparison of NIR laser light (980 nm, 1.0 W/cm^2^, 10 min) induced temperature increases in Ce6 and Ce6-LUMSN samples (33 µg/mL Ce6) in 10 mM phosphate buffer (pH 7.4). (**h**) Thermal images of saline, Ce6 and Ce6-LUMSN (33 µg/mL Ce6) samples illuminated with NIR laser light (980 nm, 1.5 W/cm^2^) for 5 min. (**g**) Photothermal stability of Ce6-LUMSNs (150 µg/mL nanospheres) monitored over five consecutive NIR laser irradiation (980 nm, 1.5 W/cm^2^, 5 min) on/off cycles. (**i**) Photothermal response profile of Ce6-LUMSNs (150 µg/mL nanospheres) subjected to 980 nm laser irradiation (1.5 W/cm^2^, 10 min) followed by natural cooling. * *P* < 0.05, *** *P* < 0.001 for comparisons with controls or amongst the different samples.

### Characterization of lipid/PEG-coated UMSNs (LUMSNs)

Nanocarriers for drug delivery applications are typically coated with lipid bilayers to improve biocompatibility, colloidal stability and controlled therapeutic cargo release^37, 38^. Lipid coatings also offer the advantage that they can be readily functionalized to achieve tissue- and cell-specific targeting^39^. Moreover, the lipid bilayer coat can be doped with an inert, water-soluble polymer, such as polyethylene glycol (PEG), that reduces aggregation and minimizes interactions with serum components that mediate the phagocytic clearance^40^.

We used a previously published protocol (see Methods) to coat the surface of Ce6-loaded UMSNs (Ce6-UMSNs) with a bilayer consisting of DPPC, cholesterol and DSPE-PEG_2000_-maleimide (Figure 1a). Contacts between the bilayer coat and UMSN are likely stabilized by van der Waals and electrostatic interactions between the phospholipid headgroups and the negatively charged UMSN. The phospholipid DPPC was chosen due to its saturated acyl chains as unsaturated lipids have been shown to reduce the long-term colloidal stability of lipid-coated mesoporous silica nanocarriers^41^. Cholesterol was used to decrease the bilayer fluidity and, in turn, reduce the baseline leakage of the Ce6 encapsulated in the pores of the UMSNs^38, 41^. Finally, PEGylated DSPE was added to increase the *in vivo* circulation half-life of the nanospheres^41, 42^, with the maleimide group on the PEG facilitating functionalization with a cancer targeting moiety. The composition of the bilayer (DPPC/cholesterol/DSPE-PEG_2000_-maleimide at a 77.5:20:2.5 molar ratio) was chosen as it was reported to yield high colloidal stability and cargo loading, as well as negligible baseline cargo leakage^37^.

Transmission electron microscopy (TEM) images showed the lipid/PEG layered over the surface of UMSNs (Figure 2c). Coating was further confirmed by DLS measurements, which showed a homogenous colloidal solution (polydispersity index = 0.11 ± 0.02)^38, 43^ of lipid/PEG-coated UMSNs (LUMSNs) that have an expectedly larger hydrodynamic diameter (180 ± 10 nm) compared to UMSNs (Figure 2g; Supporting Table 1). This translates to a lipid/PEG bilayer coat thickness of ∼10 nm. It should be noted that the discrepancy in the lipid/PEG bilayer thickness between the TEM images (Figure 2c) and DLS measurements (Figure 2g) is likely due to unavoidable differences in the experimental conditions (aqueous solution vs dehydrated sample for DLS vs TEM, respectively), and the fact that PEG is not visible by electron microscopy^44^. Additionally, the zeta potential changed from −6 to −20 mV after lipid/PLGA coating (Figure 2h; Supporting Table 2), which is in agreement with the values reported for other lipid-coated mesoporous silica nanocarriers^37^.

The colloidal stability of LUMSNs was assessed to gauge their compatibility for tumor targeting and cancer therapy applications^45, 46^. There was no change in the hydrodynamic diameter of the nanospheres in 10 mM phosphate buffer at pH 7.4 (180 ± 10 nm), sodium acetate buffer solution at pH 5.5 (180 ± 15 nm) or complete cell culture medium (RPMI 1640 containing 10% fetal bovine serum (FBS), pH 7.4) (183 ± 10 nm) over 72 h (Supporting Figure 4; Supporting Table 1). Remarkably, long-term monitoring of LUMSNs revealed that they are stable for at least a month in complete medium (Figure 2k). These results indicate that coating the nanospheres with lipid/PEG leads to colloidal stabilization and acquisition of stealth properties (i.e. prevention of serum protein adsorption), and suggests that the LUMSNs are able to maintain an appropriate size in circulation that would aid in tumor localization and internalization into cancer cells^47^.

**Figure 4.**
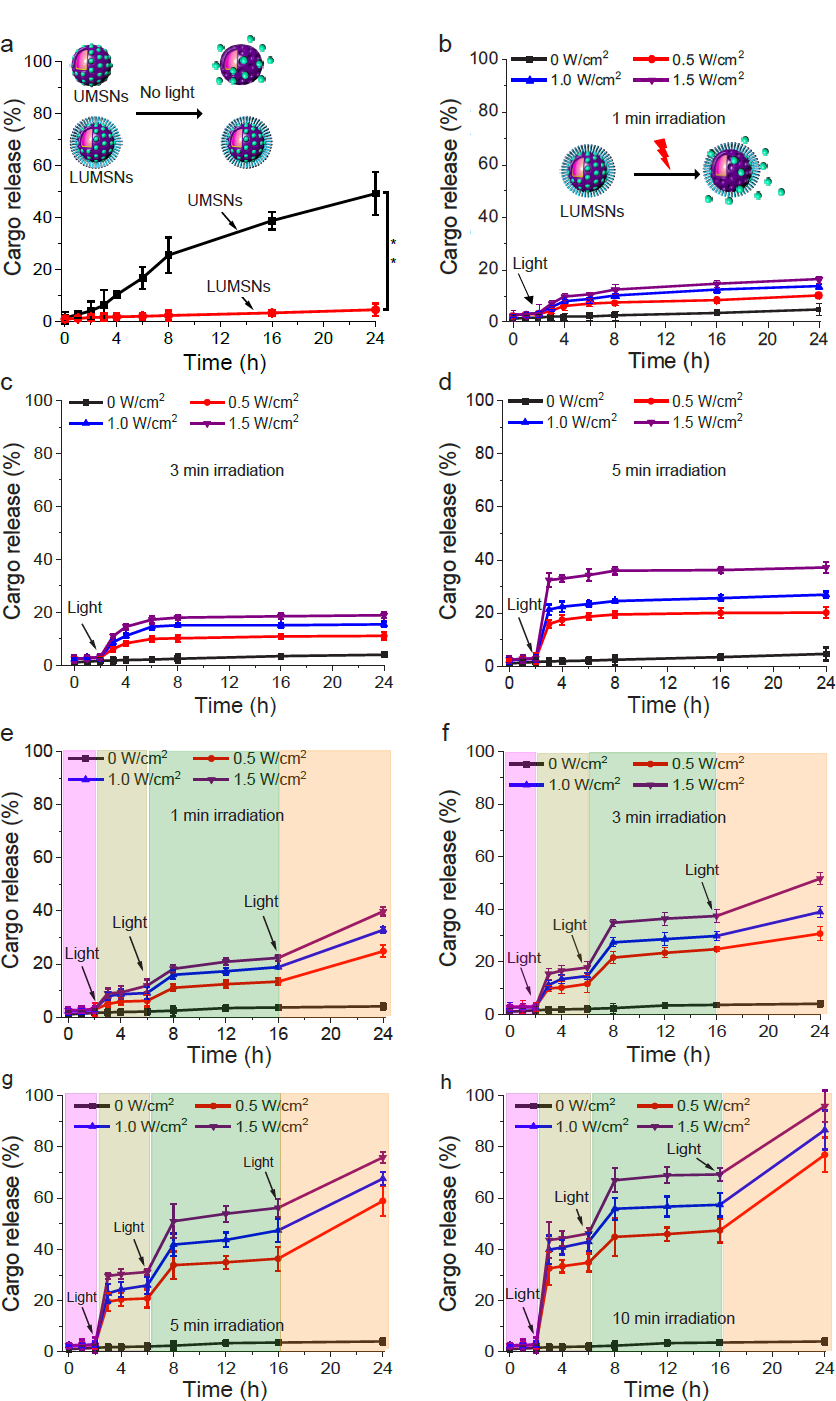
Cargo release profiles of Ce6-loaded LUMSNs in the absence and presence of a stimulus. (**a**) Release of Ce6 from UMSNs and LUMSNs in the absence of near-infrared (NIR, 980 nm) laser irradiation. (**b**–**d**) NIR laser light-triggered release of Ce6 from LUMSNs (50 µg/mL nanospheres) at varying irradiation power densities (0.5–1.5 W/cm^2^) and durations of 1 (**b**), 3 (**c**) or 5 (**d**) min in 10 mM phosphate buffer (pH 7.4). (**e**–**h**) On-demand release of Ce6 from LUMSNs (50 µg/mL nanospheres) due to sequential illumination with NIR laser light at varying irradiation power densities (0.5–1.5 W/cm^2^) and durations of 1 (**e**), 3 (**f**), 5 (**g**) or 10 (**h**) min in 10 mM phosphate buffer (pH 7.4). \*\**P* < 0.01 for comparison between UMSNs and LUMSNs.

Formation of a serum protein corona during *in vivo* circulation not only destabilizes nanocarriers and hinders their ability to target cancer cells, it also leads to immune recognition and rapid blood clearance, all of which culminates in poor tumor accumulation^23, 24^. Therefore, we investigated adsorption of serum proteins to the nanospheres further using quantitative proteomics (Supporting Figure 5; Supporting Table 3). Following incubation in complete cell culture medium (RPMI 1640 containing 10% FBS, pH 7.4) for 72 h, serum proteins that had adsorbed to the surface of UMSNs and LUMSNs were first isolated by centrifugation and then quantified using liquid chromatography tandem mass spectrometry (LC-MS/MS) with label free-quantification (LFQ)^48^. Analysis of the 144 most abundant serum proteins, selected after filtering the unavoidable contaminants, revealed markedly lower adsorption to the surface of LUMSNs compared to UMSNs (Supporting Figure 5; Supporting Table 3). Thus, the lipid/PEG coat effectively blocks the formation of a serum protein corona on the surface of LUMSNs.

**Figure 5.**
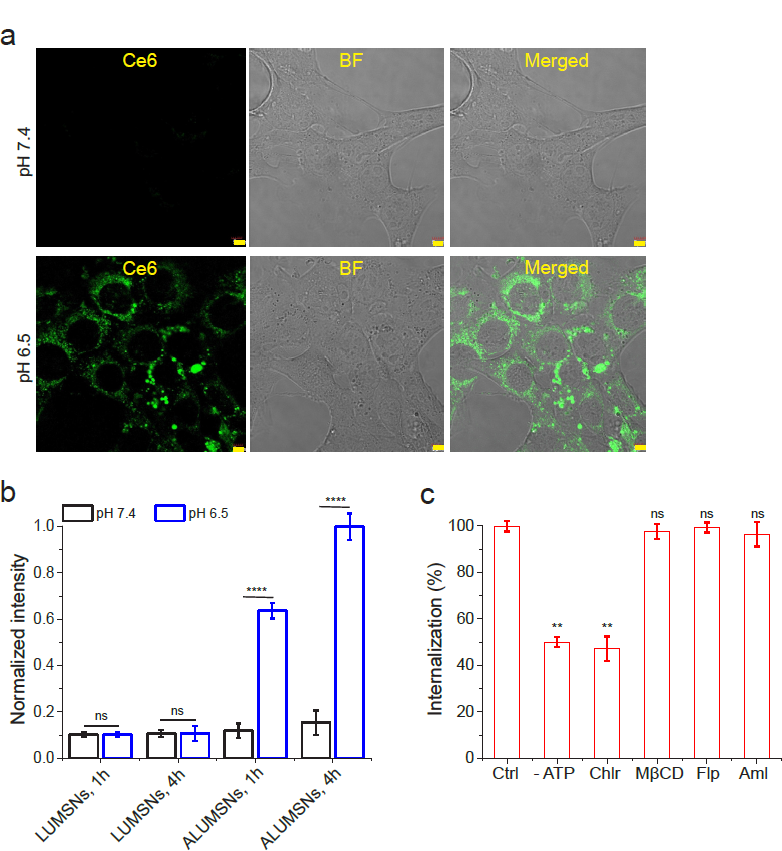
pH-dependent cellular uptake of Ce6-loaded ALUMSNs. (**a**) Confocal fluorescence microscopy images of 4T1 cells incubated with Ce6-ALUMSNs (0.5 µg/mL Ce6) for 4 h at physiological (*top panels*) or acidic (*lower panels*) pH. Ce6 is pseudo-colored green for clarity. Imaging experiments were performed in quadruplicate and representative images are shown. Scale bar = 5 µm. (**b**) Flow cytometry quantification of cellular uptake of Ce6-loaded LUMSNs and ALUMSNs (0.5 µg/mL Ce6) in 4T1 cells following treatment for 1 or 4 h at pHs 7.4 or pH 6.5 (*n* = 4). (**c**). Flow cytometry quantification of cellular uptake of Ce6-ALUMSNs (0.5 µg/mL Ce6) at pH 6.5 in 4T1 cells pretreated with sodium azide and 2-deoxy-D-glucose to deplete cellular ATP (-ATP), or with endocytosis inhibitors–chlorpromazine (Chlor; clathrin-dependent endocytosis), methyl-β-cyclodextrin (MβCD; lipid raft-mediated endocytosis), filipin (Flp; caveolae-dependent endocytosis) or amiloride (Aml; macropinocytosis inhibitor) – compared with uninhibited uptake in control cells (Ctrl) (*n* = 4). \*\**P* < 0.01, \*\*\*\**P* < 0.0001 or non-significant (ns, *P* > 0.05) for comparisons with controls or amongst the different treatment groups.

### Photodynamic and photothermal properties of Ce6-loaded LUMSNs

Ce6 is a widely used, FDA-approved second-generation PS that is characterized by high singlet oxygen (^1^O_2_) quantum yield and low dark toxicity^49–51^. However, Ce6 is prone to aggregation in solution, due to the presence of several alkyl groups, which attenuates the PS’s ^1^O_2_ production capacity^20, 50^. While chemical modification and conjugation of Ce6 to various nanocarriers has been utilized to minimize aggregation, this often decreases the PS’s ^1^O_2_ quantum yield^20^. Here, we instead loaded Ce6, without chemical modification, into the pores of LUMSNs. We monitored ROS production capability of Ce6-LUMSNs using the fluorescent probe Singlet Oxygen Sensor Green (SOSG)^52, 53^. Following NIR laser illumination, substantially higher ROS was detected in the presence of Ce6-LUMSNs compared to free Ce6, at the same Ce6 concentration and irradiation power density and duration (Figure 2l). This enhancement indicates that LUMSNs effectively prevent aggregation of the encapsulated Ce6 and the concomitant decrease in its ^1^O_2_ quantum yield.

Biocompatible and biodegradable Bi_2_Se_3_-based nanomaterials have been reported to exhibit strong NIR absorption and high photothermal conversion efficiency. We therefore investigated the temperature changes induced by NIR laser illumination of the PTA-doped LUMSNs in aqueous solution. As expected, no change in temperature was observed in Ce6-LUMSN samples in the absence of NIR light (Figure 3a–e, g). However, upon 980 laser irradiation, the Ce6-LUMSNs showed a robust, concentration and irradiation power density/duration dependent, photothermal response (Figure 3a–e, g). For instance, at 150 µg/mL LUMSNs the temperature increased from 27.1 ± 0.4 to 49.3 ± 0.3 or 53.5 ± 0.6 °C with 1.0 or 1.5 W/cm^2^ irradiation, respectively, for 5 min (Figure 3d). In contrast, a negligible increase in temperature (∼28 °C) was recorded in the free Ce6 sample compared to Ce6-LUMSNs under the same experimental conditions (Figure 3f, g), which indicates that the photothermal property of LUMSNs is due to the presence of Bi_2_Se_3_. This is supported by the photothermal response profile of the LUMSNs (Figure 3h), which matches that of various Bi_2_Se_3_-based nanomaterials^34^. Importantly, the rate of LUMSN-induced temperature increase was rapid (e.g., at a concentration of 150 μg/mL, with 1.0–1.5 W/cm^2^ irradiation, the temperature rose to ∼45–50 °C within the first 5 min; Figure 3f, h), which suggests that the nanospheres could rapidly and efficiently convert the 980 nm laser energy into heat of a temperature that is high enough to ablate the malignant cells. Additionally, the photothermal stability of LUMSNs was assessed over five laser on/off cycles. The maximum temperature (∼55 °C) was nearly identical over the five successive heating/cooling cycles, which demonstrates the high photostability of the LUMSNs (Figure 3i)^34^. Taken together, these results underline the PDT and PTT potential of the designed Ce6-LUMSNs.

### NIR light-triggered cargo release profile of LUMSNs

In addition to their photodynamic and photothermal properties, we assessed the capacity of the designed nanospheres to function as a controlled release cancer therapeutic delivery platform. In the absence of 980 nm laser irradiation, UMSNs released ∼50% of encapsulated Ce6 over the 24 h duration of measurement due to diffusion of the PS out of the pores of the uncoated nanospheres (Figure 4a). In contrast, LUMSNs did not exhibit significant stimulus-free leakage of the cargo over 24 h (Figure 4a). Thus, wrapping the nanospheres with a lipid/PEG coat resulted in a highly stable nanocarrier, which is critical for preventing premature release and ensuring that the therapeutic payload reaches the target cancer cells.

Under continuous irradiation with the 980 nm laser at varying irradiation power densities (0.5–1.5 W/cm^2^) and durations (1–5 min), the encapsulated Ce6 was efficiently released from LUMSNs (Figure 4b–d). This effect can be attributed to NIR light-induced hyperthermia increasing fluidity and permeability of the bilayer coat. The melting temperature, T_m_, of the primary phospholipid of the bilayer, DPPC, is ∼41 °C^54^, while the temperature of the LUMSNs typically rises to > 45 °C following irradiation (Figure 3d), which leads to payload release^55^. Remarkably, sequential NIR light illumination (0.5–1.5 W/cm^2^, 1–10 min) of the LUMSNs triggered repeated release of the Ce6 cargo, culminating in a maximum cumulative release of 40– 96% (Figure 4e–h). These results clearly show that LUMSNs exhibit robust NIR light-induced on-demand release of the encapsulated cargo, which highlights the potential of the designed nanospheres as a platform for combining phototherapies with chemotherapy.

### Cancer cell uptake of ATRAM-functionalized LUMSNs (ALUMSNs)

For tumor targeting, LUMSNs were functionalized with the pH-responsive acidity-triggered rational membrane (ATRAM) peptide (Figure 1)^56, 57^. ATRAM is a 34-amino acid peptide (sequence: N_t_-CGLAGLAGLLGLEGLLGLPLGLLEGLWLGLELEGN-C_t_) that interacts with cellular membranes in a pH dependent manner^26^. At physiological pH, ATRAM binds weakly and superficially to membranes in a largely unstructured conformation, while in acidic conditions the peptide inserts into lipid bilayers as a transmembrane α-helix (Figure 1b)^26, 56, 57^. Insertion of ATRAM into the membrane is driven by the increased hydrophobicity of the peptide due to protonation of its acidic glutamate residues^26^. Importantly, the membrane insertion p*K*_a_ of ATRAM is 6.5^26^, rendering the peptide ideal for targeting cancer cells in the mildly acidic (pH ∼6.5–6.8) microenvironment of solid tumors (Figure 1b, c)^58, 59^.

We previously established that ATRAM’s membrane insertion occurs via the peptide’s C-terminus^56^. Thus, LUMSNs were conjugated to ATRAM by covalently coupling the DSPE-PEG-maleimide of the lipid coat to the N-terminal cysteine of the peptide. The ATRAM-functionalized LUMSNs (ALUMSNs) were characterized using DLS and zeta potential measurements. As expected, conjugation of the peptide did not appreciably alter the hydrodynamic diameter of the nanospheres significantly (181 ± 10 nm) (Figure 2g; Supporting Table 1). However, the zeta potential at pH 7.4 increased from −20 mV to −11 mV (Figure 2h; Supporting Table 1), which confirms conjugation of ATRAM to the LUMSNs. Of relevance, the zeta potential of ALUMSNs falls within the range reported for other highly stable nanocarriers at physiological pH^57^. Lowering the pH to 6.5 increased the zeta potential of ALUMSNs to +11 mV, without adversely affecting the long-term colloidal stability of the nanospheres (Figure 2i; Supporting Figure 6; Supporting Table 1). These results strongly suggests that ALUMSNs would effectively target cancer cells within the acidic tumor microenvironment.

**Figure 6.**
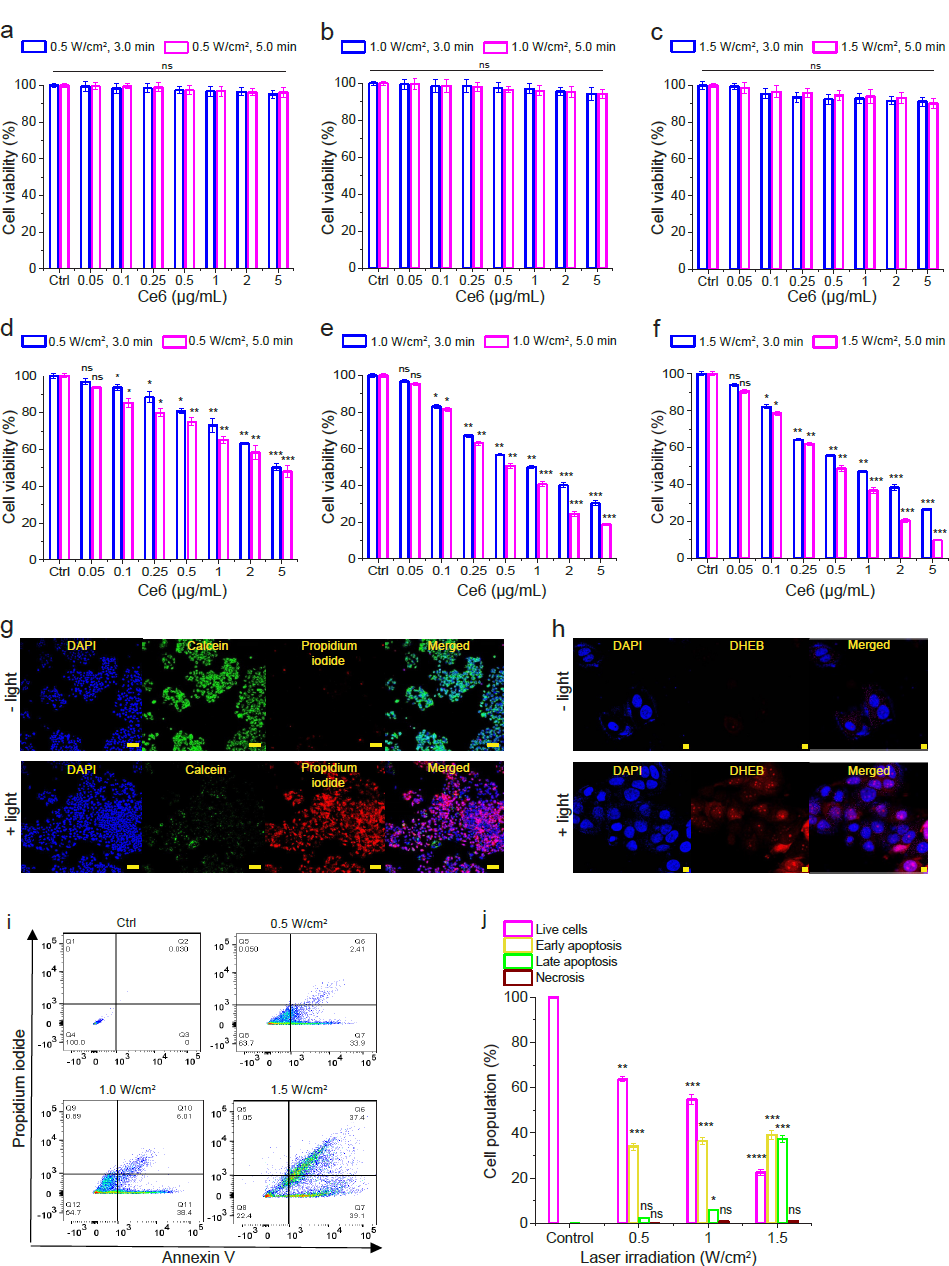
NIR light-triggered cytotoxicity of Ce6-loaded ALUMSNs. (**a**–**f**) Cell viability of 4T1 cells treated with Ce6-ALUMSNs (0.05–5 µg/mL Ce6) for 48 h and subsequently exposed to NIR (980 nm) light with different laser irradiation power densities (0.5–1.5 W/cm^2^) and durations (3.0 or 5.0 min) at pH 7.4 (**a**–**c**) or 6.5 (**d**–**f**). Cell viability in (**a**–**f**) was assessed using the MTS assay, with the % viability determined form the ratio of the absorbance of the treated cells to the control cells (*n* = 4). (**g**) Calcein AM/ propidium iodide (PI) staining of 4T1 cells incubated with Ce6-ALUMSNs (0.5 µg/mL Ce6) for 4 h in the absence (-light) or presence (+light) of NIR laser irradiation (1.5 W/cm^2^, 5 min). Scale bar = 10 μm. (**h**) Confocal fluorescence microscopy images of 4T1 cells treated with Ce6-ALUMSNs (0.5 µg/mL Ce6) for 4 h and then stained with the reactive oxygen species (ROS) probe dihydroethidium bromide (DHEB) in the absence (-light) or presence (+light) of NIR laser irradiation (1.0 W/cm^2^, 5 min). Scale bar = 10 μm. Imaging experiments in (**g**, **h**) were performed in quadruplicate and representative images are shown. (**i**) Flow cytometry analysis of annexin V/PI staining of 4T1 cells that were either untreated (control, Ctrl), or treated with Ce6-ALUMSNs (0.5 µg/mL Ce6) for 12 h at pH 6.5 and exposed to NIR light with varying laser irradiation power densities (0.5–1.5 W/cm^2^) for 5 min. The bottom left quadrant (annexin V-/PI) represents live cells; bottom right (annexin V+/PI-), early apoptotic cells; top right (annexin V+/PI+), late apoptotic cells; and top left (annexin V-/PI+), necrotic cells. (**j**) A summary of the incidence of early/late apoptosis and necrosis in the 4T1 cells treated with Ce6-ALUMSNs determined from the flow cytometry analysis of annexin V/PI staining in (**i**) (*n* = 4). \**P* < 0.05, \*\**P* < 0.01, \*\*\**P* < 0.001, \*\*\*\**P* < 0.0001 or non-significant (ns, *P* > 0.05) compared with controls.

The pH-dependent uptake of ALUMSNs in cancer cells was assessed using confocal fluorescence microscopy, TEM and flow cytometry. Murine breast cancer 4T1 cells were incubated with ALUMSNs for 4 h (Figure 5). Confocal microscopy showed substantially higher cellular internalization, and cytosolic localization, of ALUMSNs under acidic conditions compared to physiological pH (Figure 5a). Similarly, TEM revealed much greater accumulation of the nanospheres intracellularly following incubation for 4 h at pH 6.5 relative to 7.4 (Supporting Figure 7). The imaging results were confirmed with flow cytometry analysis, which showed ∼6 - and ∼9-fold higher uptake at acidic versus physiological pH at 1 and 4 h incubations, respectively (Figure 5b). In contrast, poor uptake of LUMSNs (i.e., in the absence of ATRAM) was observed in 4T1 cells at both pHs (7.4 and 6.5) and incubation times (1 and 4 h) (Figure 5b). These results confirm that ATRAM facilitates uptake of ALUMSNs specifically in cells within a mildly acidic environment.

**Figure 7.**
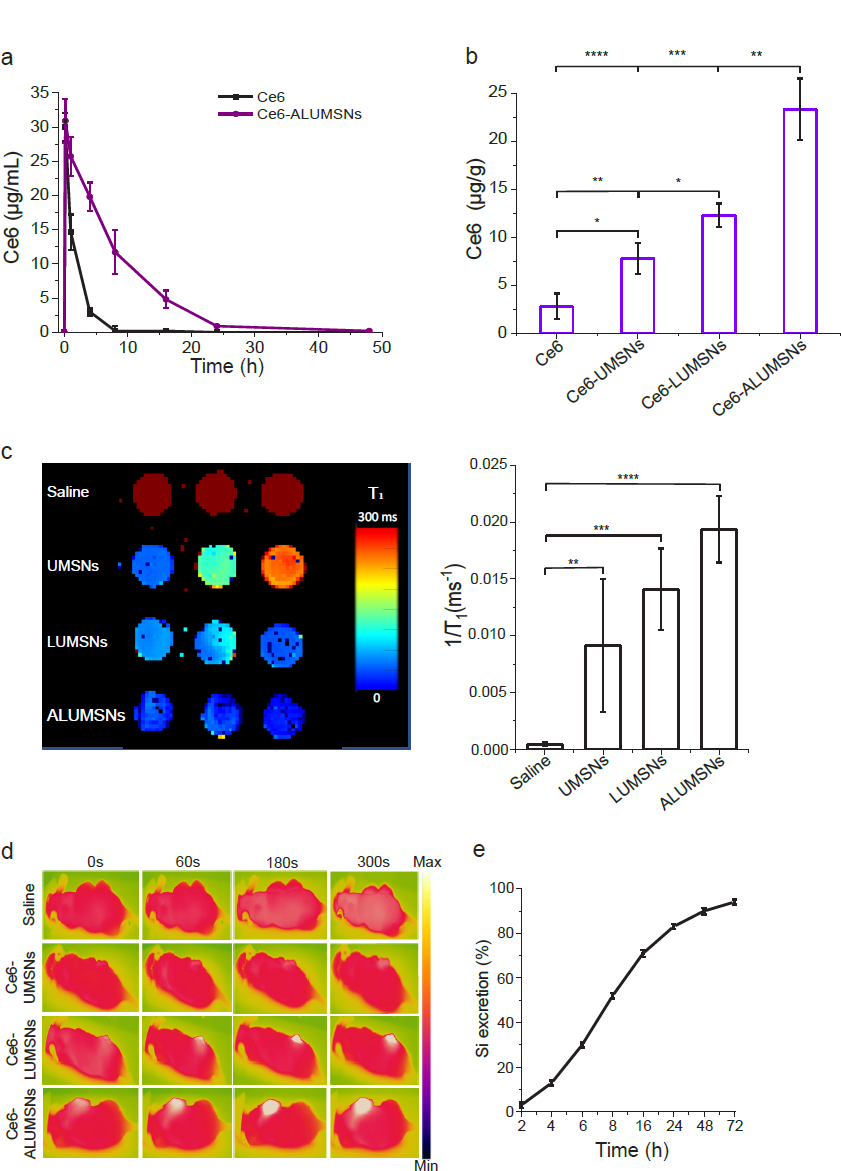
*In vivo* pharmacokinetics and biodistribution of ALUMSNs. (**a**) Concentration of Ce6 in plasma of mice following a single *i.v.* injection of free Ce6 (2.5 mg/kg) or Ce6-loaded ALUMSNs (11 mg/kg nanospheres, 2.5 mg/kg Ce6) (*n* = 4 per group). (**b**) Concentration of Ce6 in 4T1 tumors in mice 8 h after a single *i.v.* injection of free Ce6 (2.5 mg/kg) or Ce6-loaded UMSNs, LUMSNs or ALUMSNs (11 mg/kg nanospheres, 2.5 mg/kg Ce6) (*n* = 4 per group). Ce6 concentration in (**a**, **b**) was quantified using high performance liquid chromatography (HPLC)^81^. (**c**) T_1_ maps (*left*) and relaxation rates (1/T_1_) (*right*) of 4T1 tumors isolated from mice 6 h following *i.v.* injection with saline or nanospheres (UMSNs, LUMSNs or ALUMSNs; 11 mg/kg) (*n* = 3 per group). (**d**) Thermal imaging of 4T1 tumor-bearing mice upon 980 nm laser irradiation (1.0 W/cm^2^, 5 min) at different timepoints (0–300 s) post *i.v.* injection with saline or Ce6-loaded UMSNs, LUMSNs or ALUMSNs (11 mg/kg nanospheres, 2.5 mg/kg Ce6) (*n* = 4 per group). (**e**) Cumulative percentage of Si in urine and feces collected form test mice at various timepoints (2–72 h) post *i.v.* injection of ALUMSNs (11 mg/kg) (*n* = 4) determined by inductively coupled plasma mass spectrometry (ICP-MS)^42^. \**P* < 0.05, \*\**P* < 0.01, \*\*\**P* < 0.001, \*\*\*\**P* < 0.0001 or non-significant (ns, *P* > 0.05) for comparisons amongst the different treatment groups.

Next, we performed a series of experiments to elucidate the cellular internalization mechanism(s) of ALUMSNs. Depleting intracellular ATP using sodium azide/deoxyglucose markedly decreased internalization, indicating that ALUMSNs are taken up by both energy-dependent (e.g., endocytosis) and energy-independent (i.e., direct translocation) processes (Figure 5c). The direct translocation mechanism likely entails ATRAM-mediated anchoring followed by fusion of the lipid-based coat with the cancer cell membrane and concomitant release of the UMSNs into the cytosol^57^.

To determine the nature of the energy-dependent uptake mechanism, the cells were pretreated with specific endocytosis inhibitors: chlorpromazine (clathrin-coated pit formation inhibitor)^60^, methyl-β-cyclodextrin (disrupts lipid raft-mediated endocytic pathways by depleting plasma membrane cholesterol)^61^, filipin (caveolae-dependent endocytosis inhibitor)^62^, or amiloride (Na^+^/H^+^ exchange inhibitor that blocks micropinocytosis)^63^. Of all the inhibitors tested, only chlorpromazine significantly diminished cellular internalization, which indicates that clathrin-mediated endocytosis contributes to the uptake of ALUMSNs (Figure 5c). In the case of direct translocation across the plasma membrane, ALUMSNs would gain direct access to the cytosol; on the other hand, following uptake by clathrin-mediated endocytosis, acidification of mature endocytic compartments would promote endosome membrane insertion and destabilization by ATRAM, similar to other pH-responsive peptides, leading to release of ALUMSNs into the cytosol^64, 65^. Thus, the pH-dependent cellular uptake of ALUMSNs occurs by multiple mechanisms, which enables the nanospheres to efficiently internalize into tumor cells.

### Cytotoxicity of Ce6-loaded ALUMSNs

The anticancer activity of the designed nanospheres was evaluated using the MTS cell viability assay, which measures reduction of the tetrazolium compound MTS to a soluble product, formazan, by dehydrogenase enzymes in live cells^66, 67^. In the absence of NIR laser irradiation, treatment with Ce6-free UMSNs or LUMSNs (5–100 µg/mL) did not significantly reduce breast cancer 4T1 cell viability at either physiological or acidic pH (Supporting Figure 8a). Likewise, without NIR laser light the Ce6-loaded ALUMSNs (Ce6-ALUMSNs) were not toxic to 4T1 cells, up to a Ce6 concentration of 5 µg/mL, at pH 7.4 or 6.5 (Supporting Figure 8b). These results confirm that the nanospheres are biocompatible and therefore well-suited for cancer therapy applications.

**Figure 8.**
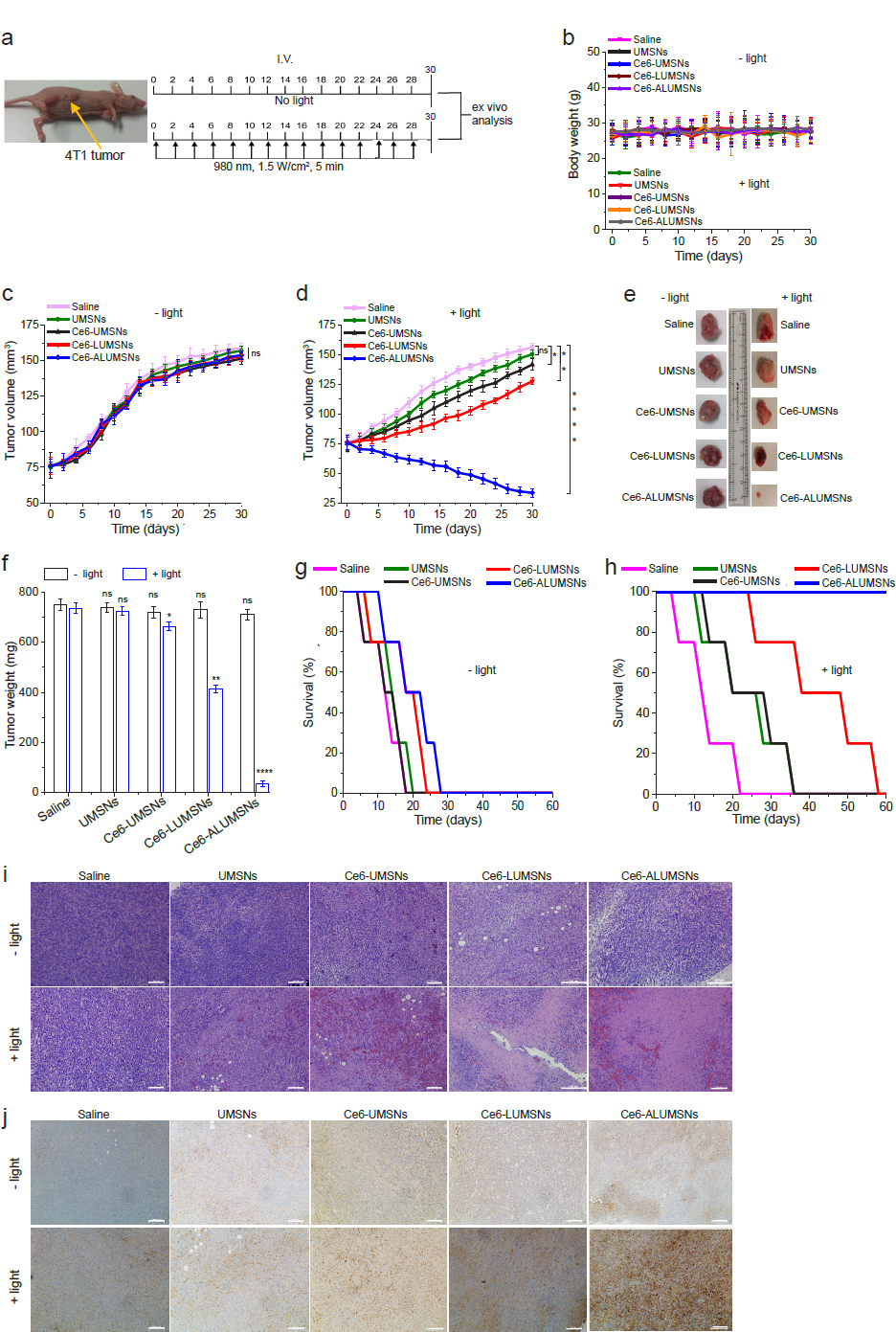
Inhibition of 4T1 tumor growth by Ce6-loaded ALUMSNs. (**a**) Design of the tumor reduction studies. Once the tumor volume reached ∼75 mm^3^, the mice were randomized into the different treatment groups (*n* = 16 per group), which were injected intravenously with saline, UMSNs (11 mg/kg) or Ce6-loaded UMSNs, LUMSNs or ALUMSNs (11 mg/kg nanospheres, 2.5 mg/kg Ce6). Injections were done every 2 days for a total of 15 doses, with the first day of treatment defined as day 0. Within each treatment group, half of the mice were subjected to NIR laser irradiation (980 nm, 1.5 W/cm^2^, 5 min) at 6 h post injection. (**b**) Bodyweight changes of the 4T1 tumor-bearing mice in the different treatment groups in the absence (-light) or presence (+light) of irradiation monitored for the duration of the experiment. (**c**, **d**) Tumor volume growth curves for the 4T1 tumors in the saline, UMSN, Ce6-UMSN, Ce6-LUMSN and Ce6-ALUMSN treatment groups over 30 days of treatment in the absence (**c**) or presence (**d**) of NIR laser irradiation (*n* = 8 per group). Tumor volume was measured via high-precision calipers using *Equation 2*. (**e**, **f**) Tumor mass analysis for the saline, UMSN, Ce6-UMSN, Ce6-LUMSN and Ce6-ALUMSN treatment groups. After 30 days of treatment, 4 mice per treatment group were sacrificed and the tumor tissues were isolated and imaged (**e**) and subsequently measured weighed to determine the tumor mass (**f**). (**g**,**h**) Survival curves for the different treatment groups (saline, UMSNs, Ce6-UMSNs, Ce6-LUMSNs and Ce6-ALUMSNs) over 30 days in the absence (**g**) or presence (**h**) of NIR laser irradiation (*n* = 4 per group). (**i**) Hematoxylin and eosin (H&E) staining of 4T1 tumor sections from the different groups (saline, UMSNs, Ce6-UMSNs, Ce6-LUMSNs and Ce6-ALUMSNs) after 30 days of treatment in the absence (-light) or presence (+light) of NIR laser irradiation. Scale bar = 500 μm. (**j**) Immunohistochemistry (IHC) images of 4T1 tumor sections stained using the caspase-3 antibody from the different groups after 30 days of treatment in the absence (-light) or presence (+light) of NIR laser irradiation. Scale bar = 1000 μm. \**P* < 0.05, \*\**P* < 0.01, \*\*\**P* < 0.001, \*\*\*\**P* < 0.0001 or non-significant (ns, *P* > 0.05) for comparisons with controls.

In the presence of 980 nm laser light, Ce6-ALUMSNs did not adversely affect 4T1 cell viability at pH 7.4 (Figure 6a–c), which is to be expected given the poor cell internalization of the nanospheres at physiological pH (Figure 5a, b). In contrast, Ce6-loaded ALUMSNs were highly toxic to the cells at pH 6.5, and the toxicity of the nanospheres scaled with PS concentration and laser power density/irradiation duration (Figure 6d–f). The MTS assay results were supported by calcein AM/propidium iodide (PI) staining, which showed that treatment of the cells with Ce6-ALUMSNs at pH 6.5 in combination with 980 laser irradiation resulted in a marked decrease in live cells (calcein signal), and a concomitant increase in dead cells (PI signal) (Figure 6g). Combined with the cell uptake experiments, the cell viability assays confirm that ATRAM mediates both the pH-dependent cancer cell uptake and the associated NIR light-induced cytotoxicity of the coupled nanospheres.

To elucidate the mechanism of cytotoxicity of Ce6-ALUMSNs, we carried out a number of complementary assays. First, we used the fluorescent ROS probe dihydroethidium bromide (DHEB) to detect intracellular ROS generation in 4T1 cells treated with the PS-loaded nanospheres at pH 6.5 and subsequently irradiated with NIR light. The bright red DHEB fluorescence signal observed in the confocal microscopy images reflects increased intracellular ROS levels upon NIR laser illumination (Figure 6h). Next, we used the fluorescent probe tetramethylrhodamine methyl ester (TMRM) to monitor mitochondrial membrane potential (ΔΨ_m_)^68^. TMRM accumulates preferentially in active mitochondria where its fluorescence intensity changes in response to changes in ΔΨ_m_^69–71^. Confocal microscopy images revealed that exposure of 4T1 cells to Ce6-loaded ALUMSNs and NIR irradiation dramatically decreased TMRM fluorescence, indicating substantial depolarization of ΔΨ_m_ (Supporting Figure 9), which agrees with reports that elevated intracellular ROS levels cause mitochondrial damage^72^. Interestingly, hyperthermia has also been shown to induce opening of the pathological mitochondrial permeability transition pore and depolarize ΔΨ_m_^73, 74^. Finally, apoptotic cells were detected using FITC-conjugated annexin V/PI staining and flow cytometry^57, 75, 76^. Treatment of 4T1 cells with Ce6-ALUMSNs at pH 6.5, followed by 980 nm laser irradiation, resulted in > 70% of the cells undergoing apoptosis (Figure 6i, j). Collectively, these results show that Ce6-ALUMSNs cause NIR light-induced toxicity selectively in cancer cells within a mildly acidic environment, and suggest that this toxicity occurs via combined ROS generation and hyperthermia that lead to ΔΨ_m_ depolarization and apoptosis.

### Macrophage recognition and immunogenicity of ALUMSNs

To prevent opsonization and subsequent uptake by monocytes and macrophages of the mononuclear phagocyte system (MPS), which can lead to undesirable accumulation in healthy tissue rather than at the target tumors^77, 78^, the nanospheres were ‘wrapped’ in a lipid/polyethylene glycol (PEG) coat^37^. PEG is a commonly used as a ‘stealth polymer’ in nanocarrier formulations to avoid opsonization and evade MPS clearance^79^.

Interaction of ALUMSNs with macrophages was assessed by first quantifying uptake of the nanospheres in differentiated human monocytic leukemia THP-1 cells, a widely used model of monocyte/macrophage activation^80^, using flow cytometry. While Ce6-loaded UMSNs were readily taken up by differentiated THP-1 cells at pH 7.4, negligible internalization of Ce6-ALUMSNs in the cells was detected under the same conditions (Supporting Figure 10a, b). Moreover, exposure to Ce6-UMSNs reduced viability of THP-1 cells, and induced production of the inflammatory cytokines tumor necrosis factor-alpha (TNF-α) and interleukin 1 beta (Il-1β) by the macrophages (Supporting Figure 10c, d). In contrast, no significant toxicity or TNF-α/Il-1β production was observed following treatment with Ce6-ALUMSNs (Supporting Figure 10c, d). These results demonstrate that ALUMSNs effectively escape recognition and uptake by macrophages, a property of the lipid/PEG-coated nanospheres that is critical for their capacity to effectively target tumors.

### Pharmacokinetics and biodistribution of ALUMSNs

4T1 tumor-bearing mice were intravenously injected with Ce6, either in free form or encapsulated in nanospheres (UMSNs, LUMSNs and ALUMSNs). Blood was then collected at specific time points and the concentration of Ce6 in the samples was measured by high performance liquid chromatography (HPLC)^81^. The *in vivo* circulation half-life of Ce6-ALUMSNs (t_1/2_ = 6.8 ± 2.1 h) was considerably longer than that of free Ce6 (t_1/2_ = 1.9 ± 0.9 h) (Figure 7a). Furthermore, while free Ce6 was eliminated from the bloodstream in ∼8 h, the PS encapsulated in ALUMSNs persisted in the plasma up to 24 h post injection. The longer *in vivo* circulation time is expected to result in greater accumulation in target tumor tissue and, in turn, increased antitumor potency^76^.

To test this hypothesis, we performed HPLC quantification of Ce6 in the 4T1 tumors, which revealed a much higher concentration of Ce6-loaded ALUMSNs (21.2 ± 5.0 μg Ce6/g of tumor tissue) compared to LUMSNs (12.5 ± 1.8 μg Ce6/g of tumor tissue), UMSNs (6.4 ± 2.5 μg Ce6/g of tumor tissue) or free Ce6 (6.3 ± 2.3 μg/g of tumor tissue) (Figure 7b). The tumor localization of the nanospheres was further investigated using MRI and thermal imaging. T_1_ mapping revealed a stronger contrast enhancement effect (i.e. lower T_1_ relaxation times) in 4T1 tumors of mice treated with ALUMSNs compared to LUMSNs and UMSNs (Figure 7c). Similarly, thermal imaging following 980 nm laser irradiation (1.0 W/cm^2^) showed that Ce6-ALUMSNs induced a much more rapid and pronounced temperature increase in the tumors (from 36 to 55 °C within 5 min) compared to other PS-loaded nanosphere formulations^82^ (Figure 7d), illustrating the *in vivo* photothermal conversion efficiency and photostability of Ce6-ALUMSNs. Of relevance, hyperthermia not only serves to ablate cancer cells, but has also been shown to increase intratumoral blood flow and enrich tumor oxygenation, which relieves tumoral microenvironment hypoxia and enhances PDT effects^17, 18, 83^. Taken together, these results suggest that the tumor-targeting ALUMSNs effectively facilitate multimodal tumor imaging (MRI and thermal imaging) and combinatorial cancer therapy (PDT and PTT).

Finally, clearance of intravenously injected Ce6-loaded ALUMSNs was determined by measuring the Si content in the urine and feces of test mice at various timepoints (2–72 h) post-injection using inductively coupled plasma mass spectrometry (ICP-MS)^42^. The advantage of ICP-MS is that it can accurately detect a wide range of elements simultaneously in a sample down to levels of ∼10 pg/mL. Consistent with other mesoporous silica-based nanoformulations^84^, most of the ALUMSNs (∼95%) were excreted via urine and feces within 72 h following administration (Figure 7e), confirming the excellent biodegradability of the nanospheres.

### *In vivo* tumor growth inhibition by Ce6-loaded ALUMSNs

Given the promising *in vitro* results of the Ce6-loaded ALUMSNs – potent and selective, NIR light-induced, anticancer activity (Figure 6) coupled with minimal interactions with serum proteins and macrophages (Supporting Figure 10) – as well as their effective tumor targeting (Figure 7), we next evaluated the antitumor efficacy of the nanospheres.

Mice bearing 4T1 mammary carcinoma tumors were injected intravenously with UMSNs (11 mg/kg) or Ce6-loaded UMSNs, LUMSNs or ALUMSNs (11 mg/kg nanospheres, 2.5 mg/kg Ce6), every 2 days for a total of 15 doses (Figure 8a). The Ce6 dose injected here is comparable to that used in other PDT-based cancer treatment studies ^85, 86^. As expected, in the absence of 980 nm laser irradiation, none of the treatments had any significant effect on growth of the 4T1 tumors (Figure 8c, e, f) or survival of the mice (Figure 8g). In the presence of NIR laser irradiation, treatment with UMSNs yielded negligible anticancer effects, which were only modestly enhanced by loading the nanospheres with Ce6 (Figure 8d–f). A more pronounced reduction in tumor growth, and a greater increase in median survival time, was observed in the Ce6-LUMSN treatment group (Figure 8d–f, h). However, treatment with Ce6-ALUMSNs yielded the greatest antitumor effects, decreasing the 4T1 tumors from an initial volume of 75 ± 7.8 to 33.5 ± 3.6 mm^3^ (Figure 8d) and the tumor mass to ∼5% that of the controls (Figure 8e, f). Ce6-ALUMSNs also prolonged survival substantially compared to the controls and all the other treatment groups over the duration of the experiment (Figure 8h). Histological (hematoxylin and eosin (H&E)) staining corroborated the greater antitumor efficacy of Ce6-ALUMSNs compared to all other treatment groups (Figure 8i). Moreover, immunohistochemistry (IHC) analysis revealed upregulation of caspase-3, a crucial mediator of apoptosis^87^, in tumor sections from the Ce6-ALUMSN treatment group (Figure 8j). This is in agreement with the *in vitro* studies, which indicated that the NIR laser light-triggered cytotoxic effects of Ce6-ALUMSNs in cancer cells is due to PDT and PTT mediated apoptosis (Figure 6).

Crucially, treatment with Ce6-ALUMSNs did not adversely affect the bodyweight of the mice (Figure 8b), and H&E staining of vital organ (lung, liver, spleen, heart and kidney) sections showed no apparent abnormalities or lesions (Supporting Figure 11), in the absence or presence of NIR laser irradiation. Taken together, these results demonstrate that the tumor targeting Ce6-loaded ALUMSNs potently shrink tumors *in vivo*, via NIR irradiation induced PDT and PTT, without adversely affecting healthy tissue, thereby markedly prolonging survival.

## CONCLUSIONS

Despite their promise as non-invasive light-based cancer treatments, PDT and PTT are currently beset by a number of issues that have hindered their clinical application. These include poor solubility, low stability, and lack of tumor specificity of many common PSs and PTAs^5, 88^. Moreover, the often-hypoxic microenvironment of tumors impairs PDT since PSs require molecular oxygen to generate ROS^88^, while hyperthermia-induced overexpression of heat shock proteins can attenuate the effects of PTT^17, 18^. Here, we have developed multifunctional core-shell nanospheres that overcome these issues. The nanospheres are composed of: a lanthanide- and PTA-doped upconversion core (NaYF_4_:Yb/Er/Gd,Bi_2_Se_3_); a PS (Ce6)-loaded mesoporous silica shell; and a lipid/PEG bilayer (DPPC/cholesterol/DSPE-PEG) coat, which is functionalized with ATRAM peptide. The ATRAM-functionalized, lipid/PEG-coated upconversion mesoporous silica nanospheres (ALUMSNs) combine the following critical properties: (i) stable encapsulation of PTAs and PSs, which prevents their aggregation and protects them from premature degradation; (ii) minimal interactions with healthy tissue, serum proteins and macrophages, leading to increased *in vivo* circulation half-life of the PTA and PS cargoes; (iii) efficient and specific internalization into cancer cells within a mildly acidic environment such as that of solid tumors; (iv) excitation by near-infrared (NIR) light, which has greater tissue penetration, lower autofluorescence and reduced phototoxicity compared to visible light; (v) MRI (due to the presence of Gd in the core) and NIR laser light-mediated thermal imaging; (vi) NIR laser light-induced PDT and PTT, the combination of which synergistically improves the efficacy of both phototherapies – PTT-induced hyperthermia increases local blood flow and leads to accumulation of molecular oxygen in tumor tissue and enhanced PDT^89^, while ROS generated during PDT can inhibit heat shock proteins in cancer cells and increase their susceptibility to PTT – resulting, in turn, in substantial antitumor effects. Taken together, our studies underline the potential of the biocompatible and biodegradable ALUMSNs as a promising nanoplatform that combines tumor targeting with multimodal diagnostic imaging and potent combinatorial therapy.

## SUPPORTING INFORMATION

Supporting Figures and Methods are available online.

## ACKNOWLEDGEMENTS

The authors thank Khulood Alawadi (Lecturer of Engineering Design, NYU Abu Dhabi) for preparing the graphic illustrations. The authors also thank the NYU Abu Dhabi Center for Genomics and Systems Biology (NYUAD-CGSB) for use of their BD FACSAria III for flow cytometry measurements. Imaging (confocal, TEM and MRI), NMR, quantitative proteomics and TEM experiments, as well as FTIR, CD and zetasizer measurements, were carried out using the Core Technology Platforms (CTP) resources at NYU Abu Dhabi. Proteomics data processing was done using the High-Performance Computing (HPC) resources at NYU Abu Dhabi. This work was supported by funding from NYU Abu Dhabi, an Al Jalila Foundation seed grant (AJF2018094) and an ADEK ASPIRE Award for Research Excellence (AARE20-371) to M.M., and from the National Institutes of Health (R35GM140846) to F.N.B.

