## Supporting Information for "pH-responsive upconversion mesoporous silica nanospheres for combined multimodal diagnostic imaging and targeted photodynamic and photothermal cancer therapy"

Short Title: ATRAM-functionalized upconversion mesoporous silica nanospheres

Keywords: ATRAM peptide; cancer therapy; mesoporous silica; magnetic resonance imaging; near-infrared light; photodynamic therapy; photothermal therapy; reactive oxygen species; tumor microenvironment; upconversion

### TABLE OF CONTENTS

#### Supporting Figures


| | |
|---|---|
| Supporting Figure 1. | Characterization of the core of upconversion mesoporous silica nanospheres (UMSNs) |
| Supporting Figure 2. | Characterization of the upconversion mesoporous silica nanospheres (UMSNs) |
| Supporting Figure 3. | Qualitative Jablonski diagram illustrating the upconversion process |
| Supporting Figure 4. | Size distribution analysis of lipid/PEG-coated UMSNs (LUMSNs) |
| Supporting Figure 5. | Quantitative proteomics analysis of serum protein adsorption to the surface of ALUMSNs |
| Supporting Figure 6. | Long-term colloidal stability of ATRAM-functionalized LUMSNs (ALUMSNs) at acidic tumoral pH |
| Supporting Figure 7. | pH-dependent cellular internalization of ALUMSNs |
| Supporting Figure 8. | Cytotoxicity of the nanospheres in the absence of NIR laser irradiation |
| Supporting Figure 9. | Mitochondrial membrane potential ($\Delta\Psi_m$) depolarization by the nanospheres |
| Supporting Figure 10. | Macrophage recognition and immunogenicity of the nanospheres |
| Supporting Figure 11. | Histological analysis of vital organs following treatment with the nanospheres |


#### Supporting Tables


| | |
|---|---|
| Supporting Table 1. | Summary of hydrodynamic diameters and zeta potentials of UMSNs, LUMSNs and ALUMSNs |
| Supporting Table 2. | Chlorin e6 (Ce6) loading capacity of the upcoversion mesoporous silica nanospheres (UMSNs) |
| Supporting Table 3. | Proteins corresponding to the UniProt Knowledgebase (UniProtKB) accession numbers shown in Supporting Figure 5 |

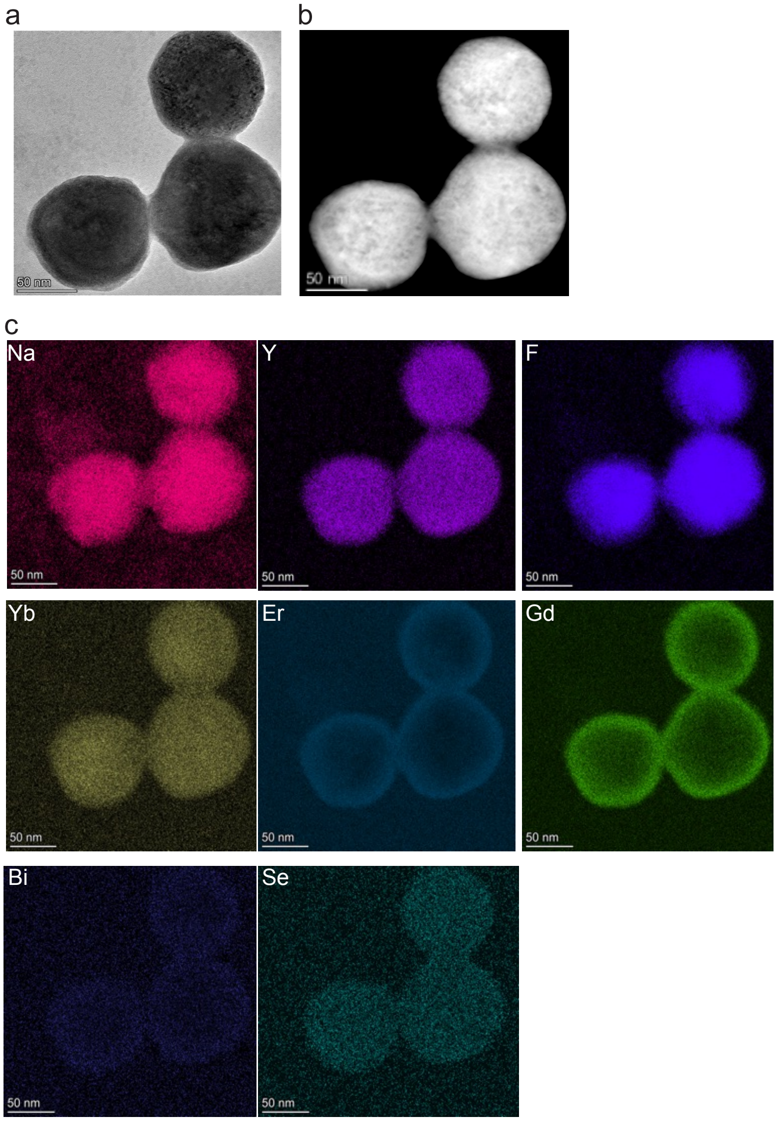


**Supporting Figure 1. Characterization of the core of upconversion mesoporous silica nanospheres (UMSNs).** (**a**,**b**) Transmission electron microscopy (TEM) (**a**) and scanning transmission electron microscopy (STEM) (**b**) images of the upconversion core ($NaYF_4$:Yb/Er/Gd,$Bi_2Se_3$) of UMSNs. (**c**) Scanning transmission electron microscopy-energy dispersive X-ray spectroscopy (STEM-EDS) mapping of the composition of the upconversion core of UMSNs. Scale bar = 50 nm.

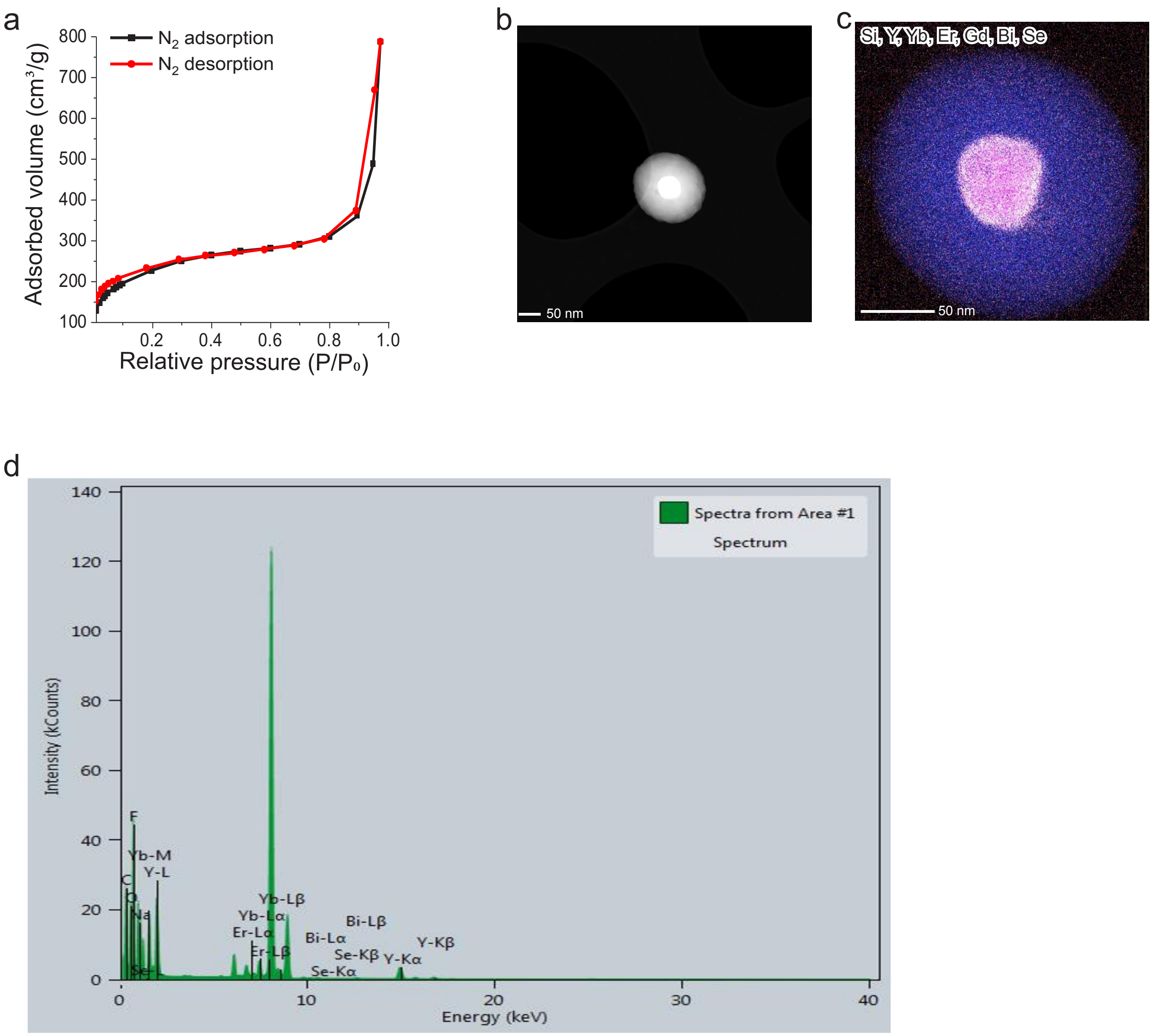


**Supporting Figure 2. Characterization of the upconversion mesoporous silica nanospheres (UMSNs).** (**a**) $N_2$ adsorption-desorption isotherms of the upconversion mesoporous silica nanospheres (UMSNs). Scale bar = 50 nm. (**b**) High-angle annular dark-field scanning transmission electron microscopy (HAAD-STEM) images of UMSNs. (**c**,**d**) Scanning transmission electron microscopy-energy dispersive X-ray spectroscopy (STEM-EDS) mapping (**c**) and the EDS spectrum (**d**) of UMSNs. Scale bar = 50 nm.

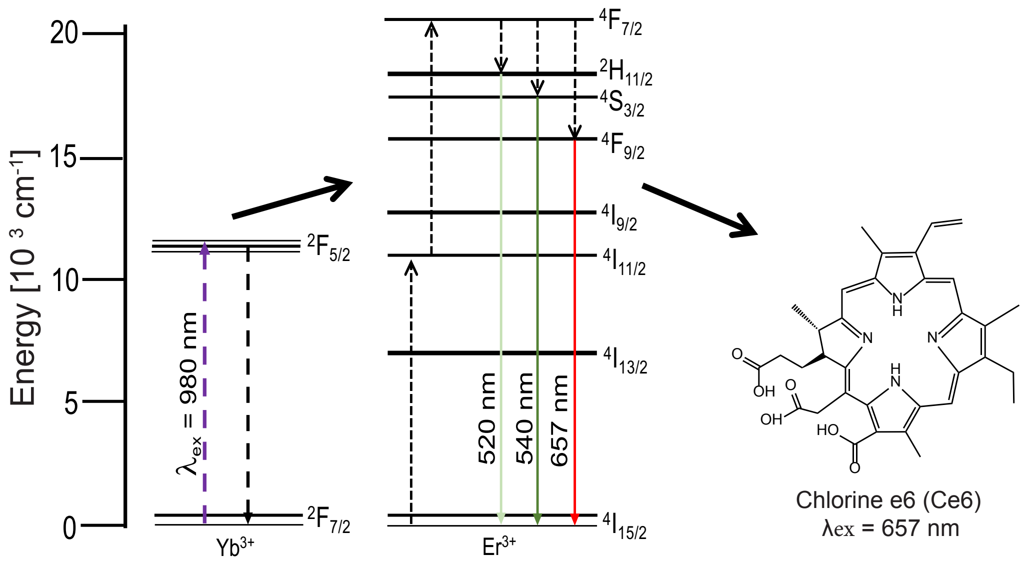


**Supporting Figure 3. Qualitative Jablonski diagram illustrating the upconversion process.** The trivalent ytterbium ion ($Yb^{3+}$) effectively absorbs near-infrared (NIR) light (980 nm) and transfers the energy to the erbium ion ($Er^{3+}$), resulting in multiple higher energy visible light emissions from $Er^{3+}$, the strongest of which are in the green and the red[1]. The red emission (657 nm) from $Er^{3+}$ excites the photosensitizer Chlorin e6 (Ce6), which then transfers its energy to molecular oxygen, converting the latter from its ground state to its cytotoxic singlet state ($^1O_2$)[1].

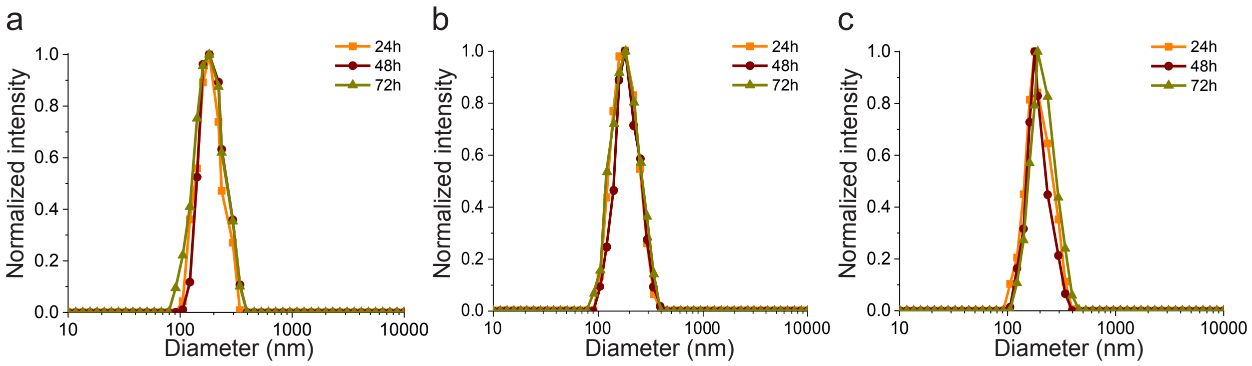


**Supporting Figure 4. Size distribution analysis of lipid/PEG-coated UMSNs (LUMSNs).** Dynamic light scattering (DLS) measurements of size of LUMSNs in 10 mM phosphate buffer (pH 7.4) (**a**), 50 mM sodium acetate buffer (pH 5.5) (**b**) and complete cell culture medium (RPMI 1640 containing 10% fetal bovine serum (FBS), pH 7.4) (**c**), over 72 h.

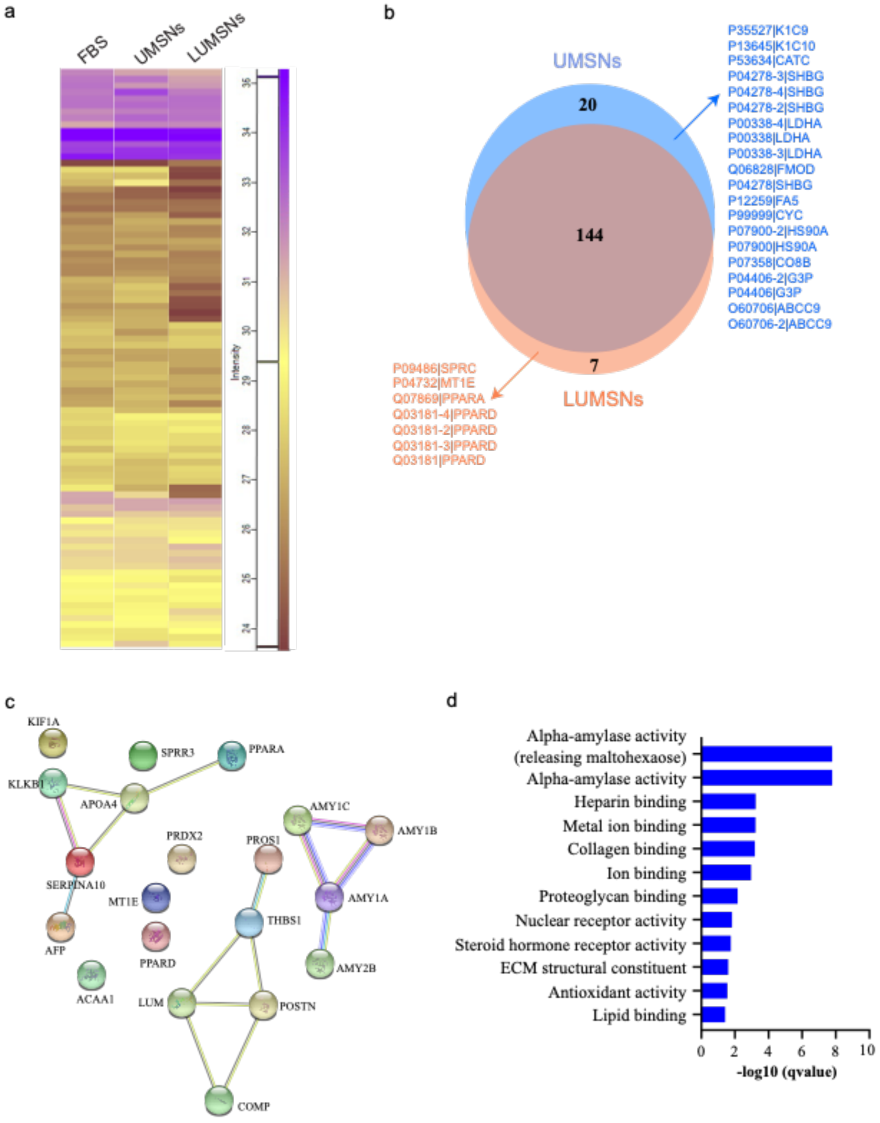


**Supporting Figure 5. Quantitative proteomics analysis of serum protein adsorption to the surface of LUMSNs.** (**a**) Heat map representation of identified serum proteins in the control fetal bovine serum (FBS) sample, and adsorbed to the surface of UMSNs and LUMSNs following incubation in complete cell culture medium (RPMI 1640 containing 10% fetal bovine serum (FBS), pH 7.4) for 72 h. The digests for both the control and nanosphere samples were analyzed by liquid chromatography tandem mass spectrometry (LC-MS/MS) and protein abundance was determined using label-free quantification (LFQ) intensities. As shown in the color scale bar, the purple and yellow (gold) colors indicate high and low LFQ intensities $(\log 2\ (LFQ))^2$ respectively, while dark brown indicates that the protein concentration is below the detection limit. The proteins corresponding to the UniProt Knowledgebase (UniProtKB) accession numbers shown in the figure are given in Supporting Table 3. (**b**) Venn diagram delineating adsorption of the 144 most abundant serum proteins to the surface of LUMSNs compared to UMSNs. (**c**) Protein–protein interaction (PPI) network map of the serum proteins depleted or absent from the surface of LUMSNs compared to UMSNs. (**d**) Gene ontology analysis for serum proteins depleted or absent from the surface of LUMSNs compared to UMSNs.

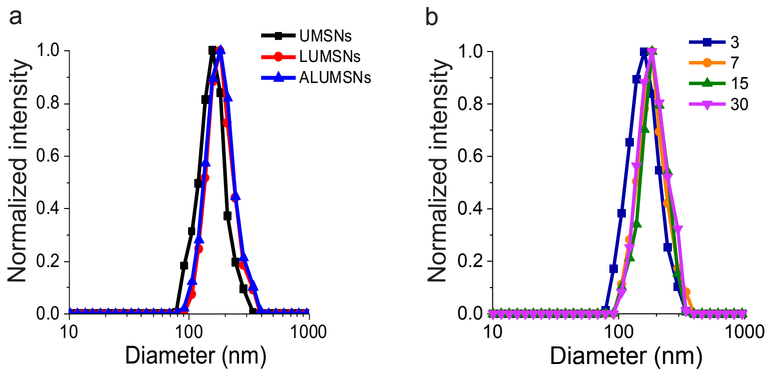


**Supporting Figure 6. Long-term colloidal stability of ATRAM-functionalized LUMSNs (ALUMSNs) at acidic tumoral pH.** (**a**) Size analysis for UMSNs, LUMSNs and ALUMSNs in complete cell culture medium (RPMI 1640 containing 10% fetal bovine serum (FBS), pH 6.5) using dynamic light scattering (DLS). (**b**) Size analysis for ALUMSNs in complete cell culture medium (RPMI 1640 containing 10% fetal bovine serum (FBS), pH 6.5) over 30 days.

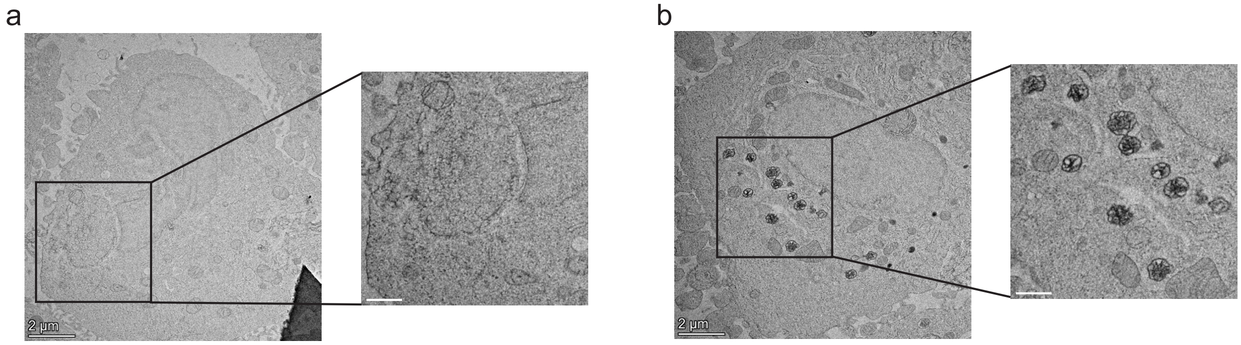


**Supporting Figure 7. pH-dependent cellular internalization of ALUMSNs.** Transmission electron microscopy images of 4T1 cells incubated with 3 µg/mL ALUMSNs for 4 h at pH 7.4 (**a**) vs 6.5 (**b**). Images on the right show magnified views of the marked areas of the images on the left. Scale bar = 2 µm.

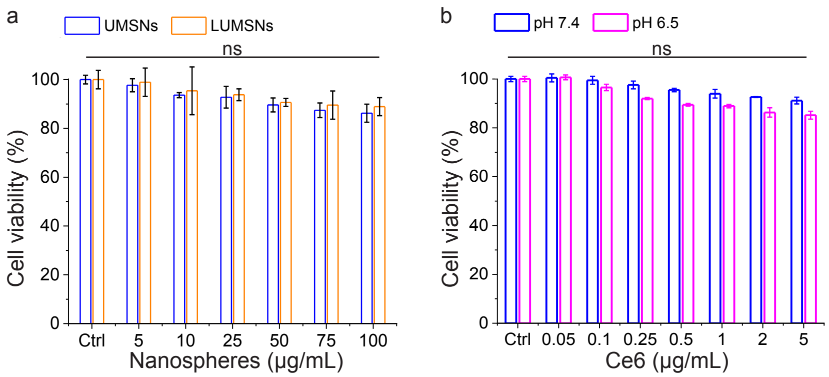


**Supporting Figure 8**. **Cytotoxicity of the nanospheres in the absence of NIR laser irradiation.** Cell viability of 4T1 cells treated with increasing concentrations of UMSNs or LUMSNs at physiological pH (**a**), or increasing concentrations of Ce6-ALUMSNs at pHs 7.4 or 6.5 (**b**). Cell viability was assessed using the MTS assay[3,4], with the % viability determined from the ratio of the absorbance of the treated cells to the control cells ($n = 4$). ns, non-significant ($P > 0.05$) for comparisons with vehicle-treated controls.

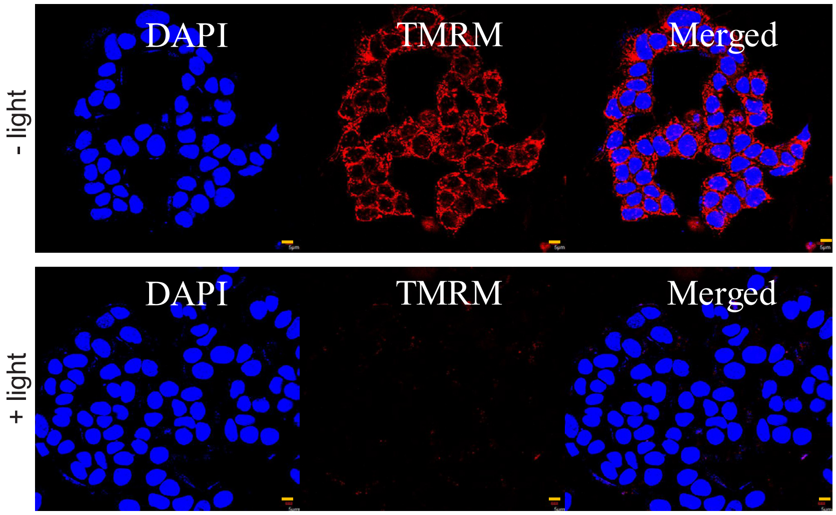


**Supporting Figure 9. Mitochondrial membrane potential ($\Delta\Psi_m$) depolarization by the nanospheres.** Confocal laser scanning microscopy images of TMRM ($\Delta\Psi_m$ reporter dye) staining of 4T1 cells treated with Ce6-loaded ALUMSNs (0.5 µg/mL Ce6) for 3 h followed by no irradiation (-light) or irradiation (+light) with NIR laser (980 nm, 1.5 W/cm$^2$, 5 min). Scale bar = 5 µm.

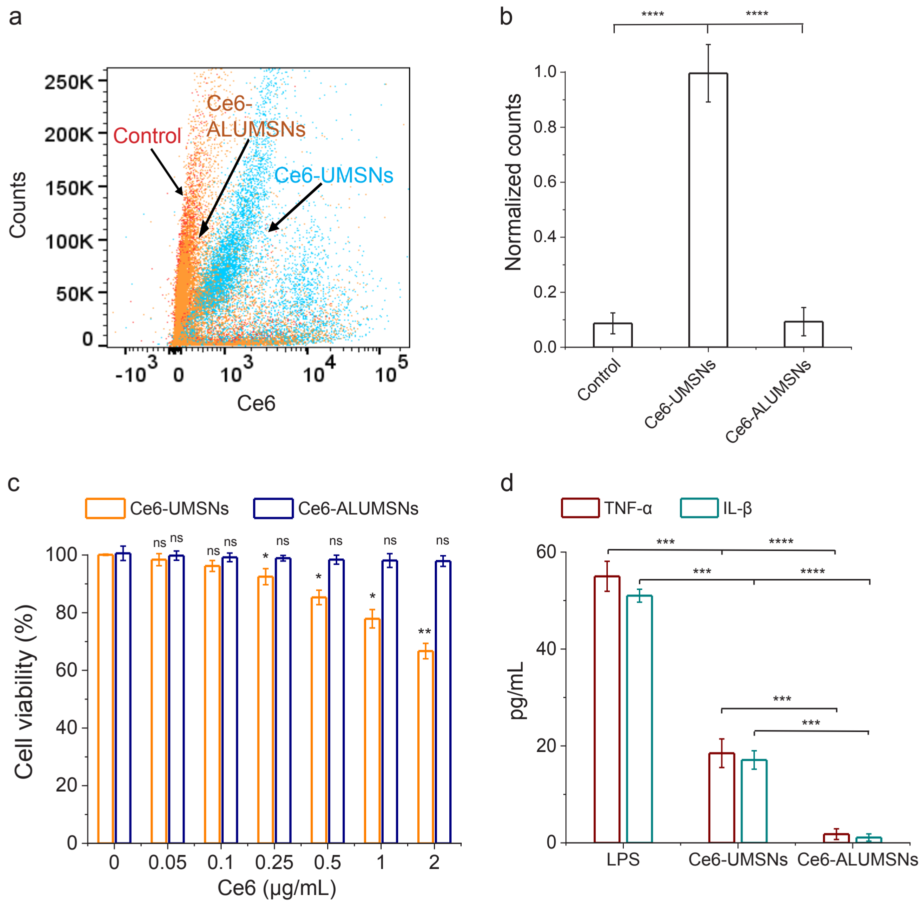


**Supporting Figure 10. Macrophage recognition and immunogenicity of the nanospheres.** (**a**,**b**) Flow cytometry analysis of differentiated THP-1 cells that were either untreated (control), or treated with Ce6-UMSNs or Ce6-ALUMSNs (0.5 μg/mL Ce6) for 4 h at pH 7.4 (**a**); quantification of cellular uptake of the nanospheres from the flow cytometry analysis ($n = 4$) (**b**). (**c**) Cell viability of differentiated THP-1 cells treated with Ce6-UMSNs or Ce6-ALUMSNs for 48 h at pH 7.4. Cell viability was assessed using the MTS assay, with the % viability determined from the ratio of the absorbance of the treated cells to the control cells ($n = 4$). (**d**) Release of inflammatory cytokines, tumor necrosis factor-alpha (TNF-α) and interleukin-1 beta (IL-1β), by differentiated THP-1 cells exposed to Ce6-UMSNs or Ce6-ALUMSNs (0.5 μg/mL Ce6) for 24 h at pH 7.4. Cells treated with lipopolysaccharide (LPS) were used as a positive control for inflammation. TNF-α and IL-1β levels in the culture medium were assayed using a commercial ELISA kit ($n = 4$). $*P < 0.05$, $**P < 0.01$, $***P < 0.001$, $****P < 0.0001$ or non-significant (ns, $P > 0.05$) compared with controls.

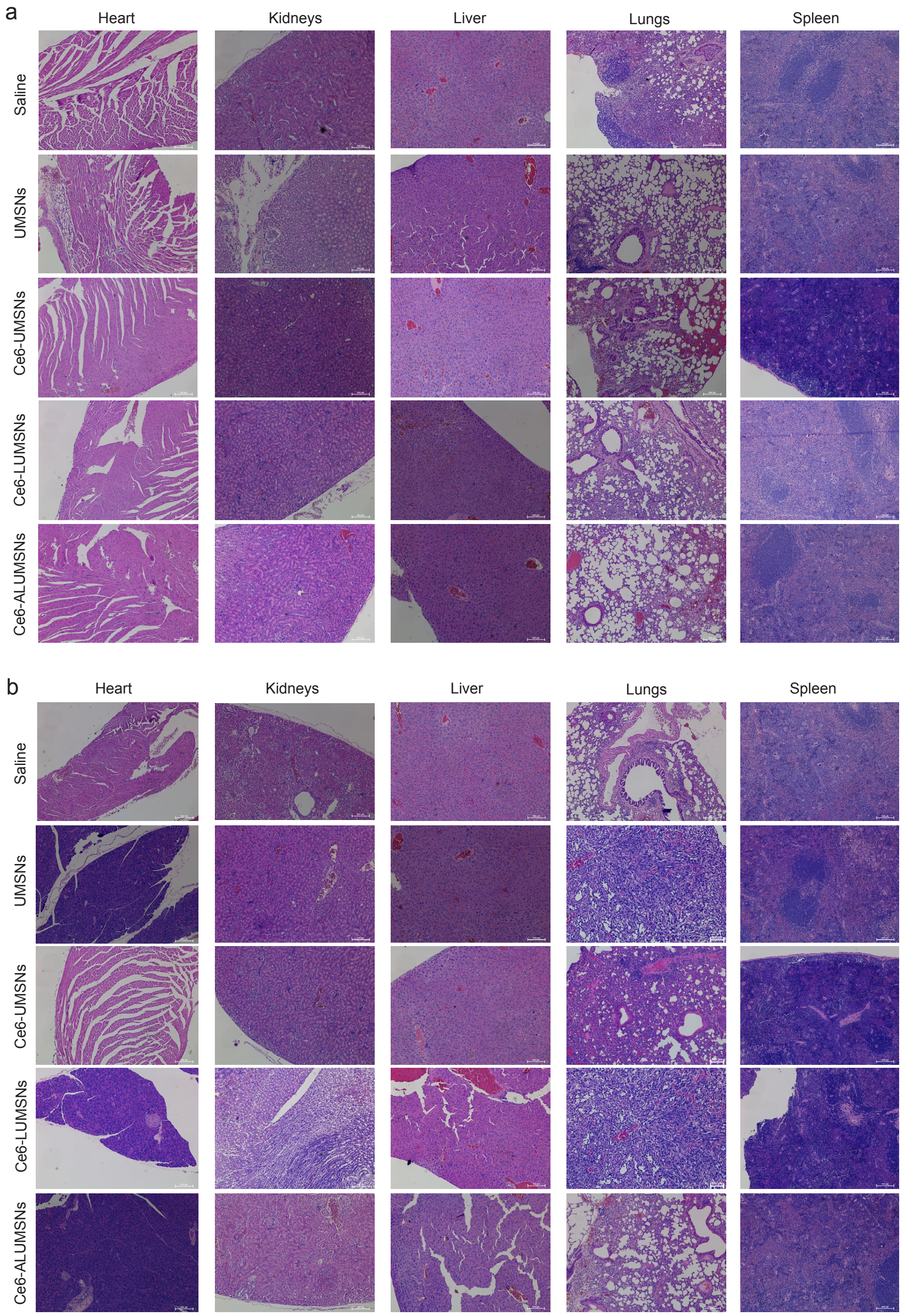


**Supporting Figure 11. Histological analysis of vital organs following treatment with the nanospheres.** Hematoxylin and eosin (H&E) staining of heart, kidney, liver, lung and spleen sections from 4T1 tumor-bearing mice after 30 days of treatment (injections administered every 2 days for a total of 15 doses) with saline, UMSNs (11 mg/kg) or Ce6-loaded UMSNs, LUMSNs or ALUMSNs (11 mg/kg nanospheres, 2.5 mg/kg Ce6), in the absence (**a**) or presence (**b**) of NIR laser irradiation (980 nm, 1.5 W/cm$^2$, 5 min) at 6 h post injection. Images shown are representative of tissue sections from four mice per treatment group. Scale bar = 50 µm.

**Supporting Table 1. Summary of hydrodynamic diameters and zeta potentials of UMSNs, LUMSNs and ALUMSNs.**

| Nanoparticle | Diameter (nm) | Zeta Potential |
|---|---|---|
| UMSNs | 160 ± 10 | -6 |
| LUMSNs | 180 ± 10 | -20 |
| ALUMSNs | 181 ± 10 | -11 (pH 7.4)<br>+11 (pH 6.5) |

**Supporting Table 2. Chlorin e6 (Ce6) loading capacity of the upcoversion mesoporous silica nanospheres (UMSNs).**

| **Ce6 feed ratio (to 5 mg UMSNs)** | **Loading capacity (wt%)** |
|---|---|
| 0.3 | 5.0 |
| 0.5 | 10.0 |
| 1.0 | 15.0 |
| 1.5 | 19.0 |
| 2.0 | 22.0 |

The Ce6 loading capacity of UMSNs was determined using *Equation 1*.

Supporting Table 3. Proteins corresponding to the UniProt Knowledgebase (UniProtKB) accession numbers shown in Supporting Figure 5.

| UniProtKB Accession Number | Protein |
|---|---|
| P09486 | SPRC |
| P04732 | Metallothionein-1E |
| Q07869 | Peroxisome proliferator-activated receptor alpha |
| Q03181 | Peroxisome proliferator-activated receptor delta |
| P35527 | Keratin, type I cytoskeletal 9 |
| P13645 | Keratin, type I cytoskeletal 10 |
| P53634 | Dipeptidyl peptidase 1 |
| P04278 | Sex hormone-binding globulin |
| P00338 | L-lactate dehydrogenase A chain |
| Q06828 | Fibromodulin |
| P12259 | Coagulation factor V |
| P99999 | Cytochrome c |
| P07900 | Heat shock protein HSP 90-alpha |
| P07358 | Complement component C8 beta chain |
| P04406 | Glyceraldehyde-3-phosphate dehydrogenase |
| O60706 | ATP-binding cassette sub-family C member 9 |

### METHODS

**Reagents.** 2-Aminoethyl dihydrogen phosphate (AEP), ammonia, ammonium bicarbonate buffer, ammonium fluoride, dead cell apoptosis kit, enzyme-linked immunosorbent assay (ELISA) kits (for detection of tumor necrosis factor - alpha human (TNF-α) and interleukin-1 beta (Il-1β)), formic acid, methanol, N-2-hydroxyethylpiperazine-N-2-ethane sulfonic acid (HEPES), octadecane (99%) and Singlet Oxygen Senor Green (SOSG), were obtained from Thermo Fisher (Waltham, MA, USA). 1,2-Dipalmitoyl-sn-glycero-3-phosphocholine (DPPC), cholesterol, and the PEGylated derivative of 1,2-distearoyl-sn-glycero-PE (DSPE-$PEG_{2000}$)-maleimide were purchased from Avanti Polar Lipids Inc (Alabaster, AL, USA). Acetone, acetonitrile, ammonium fluoride (NaF, 99.99%), bismuth(III) nitrate pentahydrate ($Bi(NO_3)_3$•$5H_2O$), calcium chloride, chloroform, cetyl trimethylammonium bromide (CTAB), cyclohexane, Dulbecco's modified eagle's MEM medium (DMEM), dimethyl sulfoxide (DMSO), dithiothreitol (DTT), ethanol, erbium(III) chloride hexahydrate ($ErCl_3$•$6H_2O$), endocytosis inhibitors (amiloride, chlorpromazine, cytochalasin D and filipin), fetal bovine serum (FBS), gadolinium(III) chloride hexahydrate ($GdCl_3$•$6H_2O$), hydrochloric acid (HCl), igepal CO-520, indole acetic acid (IAA), K3-EDTA, mannitol, octadecyl trimethoxysilane (C18-TMS), oleic acid (90%), penicillin-streptomycin, phosphate-buffered saline (PBS), polyvinylpyrrolidone (PVP), phorbol 12-myristate 13-acetate (PMA), RPMI 1640, selenium, sodium chloride (NaCl), sodium hydroxide (NaOH), sodium borohydride ($NaBH_4$), tetraethyl orthosilicate (TEOS), tetramethylrhodamine methyl ester (TMRM), trypsin, Tween 20, yttrium(III) chloride hexahydrate ($YCl_3$•$6H_2O$), and ytterbium(III) chloride hexahydrate ($YbCl_3$•$6H_2O$) were all purchased from Sigma Aldrich (St. Louis, MO, USA). Dihydroethidium (DHE) Assay kit was obtained from Abcam (Dallas, TX, USA). Gadolinium-diethylenetriamine penta-acetic acid (Gd-DTPA) was obtained from Schering AG (Berlin, Germany). The Caspase-3 Antibody 9662 was obtained from Cell Signaling Technology (Danvers, MA, USA). The CellTiter 96 Aqueous One Solution (MTS) Cell Proliferation Assay kit was from Promega (Madison, WI, USA).

**Synthesis of the upconversion core**. Initially, the upconversion hydrophobic core of the nanospheres, which consist of sodium yttrium fluoride doped with lanthanides (ytterbium, erbium, and gadolinium) and bismuth selenide ($NaYF_4$:Yb/Er/Gd,$Bi_2Se_3$) were synthesized using a thermal decomposition method with oleic acid as a capping agent[5]. Briefly, $YCl_3$•$6H_2O$ (0.99 mmol), $YbCl_3$•$6H_2O$ (0.34 mmol), $ErCl_3$•$6H_2O$ (0.06 mmol), and $GdCl_3$•$6H_2O$ (0.61 mmol) were dissolved in a round-bottom flask with a mixture of oleic acid (13.4 g) and octadecene (35 mL) under stirring and Ar atmosphere. The solution was then evacuated for 45 min until the gas evolution stopped, and the reaction mixture was heated to 140 °C under Ar atmosphere to obtain a yellow-colored solution. The solution was cooled to 45 °C, and ammonium fluoride (300 mg) and sodium hydroxide (150 mg) were added to the reaction mixture under stirring until a clear solution was obtained. The solution was heated to 300 °C and maintained for 90 min with a heating mantle, resulting in discoloration (yellow/brown) of the reaction mixture and white precipitation. The nanospheres were then separated by centrifugation (4000×*g*, 25 min) and purified with multiple redispersion and centrifugation steps in absolute ethanol. The white powder obtained was dissolved in cyclohexane after reaching room temperature.

The obtained hydrophobic cores were transferred to the aqueous phase for doping with bismuth selenide ($Bi_2Se_3$). The hydrophobic oleic acid from the surface of core nanospheres was removed using a previously reported acid-induced technique[6]. The dried core nanospheres were then redispersed in a mixture of HCl (2 M) and absolute ethanol (1:1, v/v) and sonicated (40 kHz) for 5 h at 55 °C. This process resulted in the production of oleic acid-free nanospheres, which were subsequently collected by centrifugation (4000×*g*, 25 min). After multiple rinses with ethanol, the nanospheres were dispersed in water to facilitate subsequent doping with $Bi_2Se_3$.

In order to prepare the $Bi_2Se_3$, the selenium precursor was initially synthesized by reducing 3.16 g (40 mmol) of selenium powder using 4.54 g (120 mmol) of $NaBH_4$ that was dissolved in 400 mL of aqueous solution, under an $N_2$ atmosphere at ambient temperature[7]. Then, 1 g of Mannitol was dissolved in 10 mL of water, followed by sequential addition of an aqueous solution of oleic acid free core nanospheres (2 mL, 0.125 mM), ($Bi(NO_3)_3$•$5H_2O$) (0.1g), and PVP (0.1g). The mixture was then homogeneously mixed, and the pre-synthesized selenium precursor (0.3 mmol) was added. Following 8 h of continuous stirring, the $Bi_2Se_3$-doped core nanospheres were collected by centrifugation (4000×*g*, 25 min), washed with water, and dispersed in deionized water for further use.

**Synthesis of Ce6-loaded lipid/PEG-coated upconversion mesoporous silica nanospheres (Ce6-LUMSNs).** The coating of the upconversion core to yield upconversion mesoporous silica nanospheres (UMSNs) was carried out using a previously described coating technique[8]. Briefly, 50 mL of cyclohexane, igepal CO-520 (0.8 mL), and the core nanosphere solution (in 50 mL of cyclohexane, 0.04 M) was added to a 250 mL round neck flask. Then, 1.0 mL of ammonia (28 wt%) was added, which was then sonicated for ~30 min until a transparent emulsion was formed. Following this, 200 µL of TEOS and 80 µL of C18 TMS were added, and the solution was continuously stirred for 48 h at room temperature. The obtained UMSNs were collected by centrifugation (4000×*g*, 10 min), precipitated with acetone and then washed thrice with absolute ethanol and dried at 60 °C for 12 h. Next, the synthesized UMSNs were refluxed in a mixture of methanol and HCl to remove the surfactants, and finally vacuum dried[9].

Ce6 was loaded into the UMSNs using a previously described technique for loading mesoporous silica with drugs[10]. About 5 mg of UMSNs were dispersed in 0.8 mL of DMSO and sonicated until a uniform colloidal solution was obtained. 0.2 mL of varying concentrations of Ce6 dissolved in DMSO (Supporting Table 1) were added to this solution (final volume, 1.0 mL) and allowed to stir for 48 h at room temperature. The dispersion was then centrifuged (8000×*g,* 10 min) to collect the Ce6 loaded UMSNs (Ce6-UMSNs), and the supernatant was collected to calculate the drug loading capacity. The Ce6-UMSNs were vacuum dried, washed thrice with milli-Q water to remove any residual DMSO, and lyophilized for further use. The total mass of Ce6 that was successfully loaded into the UMSNs was calculated by subtracting the mass of residual Ce6 in the isolated supernatant from the initial mass of the photosensitizer used in the nanosphere loading step. The aforementioned masses were obtained using UV-Vis spectroscopy[11], by calibrating against the standard concentrations of Ce6 (0.000313−1.0 mg/mL) in DMSO. The loading capacity of Ce6 into the UMSNs was calculated using the following equation[12]:

$$\text{Loading capacity (\%)} = ((M_0 - M_s)/W_0)\times 100 \qquad (\textit{Equation 1})$$

where $M_0$ and $M_s$ represent the initial mass of Ce6 and mass of Ce6 loaded in the UMSNs, respectively, and $W_0$ represents the mass of the UMSNs.

To wrap the nanospheres with a lipid/PEG-bilayer, 10 mg of lyophilized Ce6-UMSNs were suspended in 3 mL of saline (0.9 % NaCl) and sonicated (40 kHz) for 30 s. Then, the suspension was immediately added on top of the dehydrated lipid/PEG-film that consisted of DPPC/cholesterol/DSPE-$PEG_{2000}$ at a molar ratio of 77.5:20:2.5[10]. The Ce6-UMSNs to lipid ratio was set at 1:1.2 (w/w). Following this, probe sonication was performed for 20 min by setting the cycle at 15/15 s on/off at a power output of 30 W[10]. Then, Ce6-loaded lipid/PEG-coated UMSNs (Ce6-LUMSNs) were separated from free-lipids by centrifugation (10,000×*g*, 10 min), followed by washing twice with saline and water. Similarly, Ce6 free LUMSNs were prepared by dispersing the 10 mg UMSNs in a 3 mL of saline solution and adding them on top of the dehydrated lipid/PEG-film that consisted of DPPC/cholesterol/DSPE-$PEG_{2000}$ at a molar ratio of 77.5:20:2.5. Then probe sonication was performed for 20 min by setting the cycle at 15/15 s on/off at a power output of 30 W. Finally, Ce6 free LUMSNs were separated by centrifugation (10,000×*g*, 10 min), washed twice with saline and water and stored for further use.

**Synthesis of ATRAM-functionalized LUMSNs (ALUMSNs).** The targeting acidity-triggered rational membrane (ATRAM) peptide was synthesized by Selleck Chemicals (Houston, TX, USA) using standard Fmoc methods. The peptide was purified in house by reverse-phase high-performance liquid chromatography (Waters 2535 QGM HPLC) and its purity was subsequently verified using liquid chromatography mass spectrometry (Agilent 6538 QToF LC/MS).

In order to functionalize the LUMSNs with the ATRAM peptide, DSPE-$PEG_{2000}$-ATRAM was first synthesized by covalently conjugating DSPE-$PEG_{2000}$-maleimide with the single cysteine residue at the N-terminus[13] of ATRAM. Briefly, 600 nmol of ATRAM peptide was mixed with 500 nmol of DSPE-$PEG_{2000}$-maleimide in 200 µL of methanol, stirred overnight at room temperature under $N_2$ atmosphere, and then the conjugation was confirmed by mass spectrometry.

The dehydrated lipid film was first synthesized by mixing DPPC/cholesterol/DSPE-$PEG_{2000}$-ATRAM at a molar ratio of 77.5:20:2.5. 10 mg Ce6-UMSNs were suspended in 3 mL of saline (0.9 % NaCl) and sonicated (40 kHz) for 30 s. The suspension was then immediately added on top of the dehydrated lipid film where the Ce6-UMSNs to lipid ratio was set to 1:1.2 (w/w). They are next subjected to probe sonication for 20 min with a power output of 30 W and a cycle of 15/15 s on/off. At last, the Ce6-loaded ATRAM-functionalized LUMSNs (Ce6-ALUMSNs) and free lipids were separated by centrifugation (10,000×*g*, 10 min), followed by washing twice in saline and water.

**Characterization of the nanospheres**. All nanospheres employed in this study were thoroughly characterized to ensure their quality and validate their properties. The shape and structure of the synthesized nanospheres were characterized using transmission electron microscopy (TEM) on a Talos F200X (Thermo Fischer). To prepare the samples for TEM imaging, a drop of the respective nanosphere suspension was placed onto a 300-mesh copper TEM grid (Ted Pella, CA, USA) and dried at room temperature for 2 h to create microfilms. This process allowed us to observe the nanospheres in their dried state and obtain images with a high level of detail. The porous and textural properties of the nanospheres were analyzed using a nitrogen sorption analyzer (Micrometrics-3Flux) operated at -196 °C. The specific surface area of the samples was calculated using the Brunauer-Emmett-Teller (BET) equation. The size and zeta potential measurements were carried out using a Zetasizer Nano (Malvern Instruments; Malvern, UK) with the nanospheres suspended in different media, including water, PBS (10 mM, pH 7.4), sodium acetate buffer (50 mM, pH 5.5) and cell culture media containing 10% FBS at a particle concentration of 100 µg/mL[9].

**$T_1$ relaxation measurements**. $T_1$ relaxation quantification was performed using a 3.0 T MRI scanner (Magnetom Prisma; Siemens Healthineers, Germany) for the commercial Gd-DTPA and UMSNs at various concentrations of Gd (0.0215–0.9526 mM) in 10 mM phosphate buffer (pH 7.4). A saturation-recovery spin echo pulse sequence was employed with a range of ten repetition times (TR) from 150 ms to 10s, utilizing the following scanning parameters: echo time (TE) of 11ms, number of averages of 2, field-of-view of 180 mm × 180 mm, acquisition matrix of 320 ×

320, in-plane resolution of 0.56 mm × 0.56 mm, 20 slices, and a slice thickness of 3.5 mm. The $T_1$ values (measured in milliseconds) were quantified on a pixel-by-pixel basis from both Gd-DTPA and UMSNs at the same Gd concentration using an in-house non-linear least-squared fitting script based on the MR relaxation equation implemented in Matlab 2022. The $1/T_1$ relaxation rate was subsequently calculated as the reciprocal of $T_1$[14].

**Detection of reactive oxygen species generation**. Reactive oxygen species (ROS) generation capability of Ce6 and Ce6-LUMSNs at the same Ce6 concentration (0.5 µg/mL) in presence of near infrared (NIR) laser irradiation (980 nm, 1.0 W/cm$^2$, 5 min) was monitored in 10 mM phosphate buffer (pH 7.4) using the fluorescent probe SOSG[15] as per the previously described technique[16]. The excitation of SOSG at 490 nm with monitoring fluorescence at 550 nm allowed the assessment of the reaction of SOSG with $^1O_2$.

**Photothermal performance of nanospheres.** The temperature increase following the NIR laser irradiations (980 nm, 0–1.5 W/cm$^2$, 5 min) of 10–200 µg/mL Ce6-LUMSNs in 10 mM phosphate buffer (pH 7.4) was evaluated. Comparison of NIR laser light induced temperature increase in saline, for Ce6 and Ce6-LUMSNs samples (33 µg/mL Ce6) in 10 mM phosphate buffer (pH 7.4) was evaluated with NIR laser irradiation (980 nm, 1.5 W/cm$^2$, 5 min) and simultaneously, thermal images of the respective samples were captured. Then, photothermal stability of 150 µg/mL Ce6-LUMSNs was monitored over 5 consecutive NIR laser irradiation (980 nm, 1.5 W/cm$^2$, 5 min) on/off cycles using a previously described technique[17]. The photothermal response profile of 150 µg/mL Ce6-LUMSNs was recorded, when subjected to NIR laser irradiation (980 nm, 1.5 W/cm$^2$, 10 min) followed by natural cooling. We used an infrared camera (Optris PI-640, Optris GmBH, Germany) to record the temperature of the nanospheres, Ce6, and saline solutions.

**NIR light-triggered cargo release profile of LUMSNs.** The Ce6 release from 50 µg/mL Ce6-UMSNs and -LUMSNs was studied in triplicates using a previously described release profile technique[18] in the absence of NIR laser light irradiation. Briefly, Ce6-UMSNs and Ce6-LUMSNs were loaded into a dialysis bag (Slide-A-Lyzer MINI Dialysis Device, 2000K MWCO, 0.5 mL; Thermo Fisher). The dialysis bag was submerged into 14 mL of PBS (pH 7.4, 10 mM) containing 0.1 wt% Tween 20 and positioned on a magnetic stirrer rotating at 75 rpm. At 0–24 h, 0.5 mL of the PBS (dissolution medium) was withdrawn and replaced with 0.5 mL of fresh PBS. The obtained samples were centrifuged (1000×*g*, 5 min) and the obtained pellet containing the released Ce6 was subsequently dissolved with acetonitrile and assayed by high-performance liquid chromatography (HPLC)[19]. To measure the NIR light-triggered release of Ce6 from Ce6-LUMSNs as well any stimulus-free Ce6 leakage in the absence of irradiation, the following experimental groups were assigned:(a) Ce6-LUMSNs at varying NIR laser light irradiations (980 nm, 0.5–1.5 W/cm$^2$, 10 min), at 3 h of release experiment, (b) Ce6-LUMSNs at varying NIR laser light irradiations (0.5–1.5 W/cm$^2$) for 3.0 min, at 3 h of release experiment, (c) Ce6-LUMSNs at varying NIR laser light irradiations (980 nm, 0.5–1.5 W/cm$^2$, 5 min), at 3 h of release experiment and (d) Ce6-LUMSNs at varying NIR laser light irradiations (980 nm, 0.5–1.5 W/cm$^2$, 1–10 min) at 3, 6 and 16 h of release experiment. The amount of Ce6 released from the LUMSNs was determined following the HPLC technique[19].

**Quantitative proteomic analysis.** To investigate the interaction of serum proteins with UMSNs and LUMSNs, a solution of 1 mg/mL of the nanospheres was incubated in cell culture medium containing 10% FBS for 72 h. Following this, the nanospheres absorbed serum proteins that had adsorbed to the nanospheres were isolated through centrifugation using a described previously [20,21]. FBS was used as a control. The isolated serum proteins were then protease digested and analyzed using mass spectrometry (MS) to determine the specific proteins that had adsorbed to the nanospheres[22]. Briefly, the isolated serum proteins were reduced (using 10 mM DTT at 85 °C for 30 min) and alkylated (using 25 mM IAA for 1 h at room temperature and in the dark). The protease-specific pH was achieved by diluting the sample with 50 mM ammonium bicarbonate buffer, using a spin filter with a 30-kDa cutoff. The digests were then treated with a 1:50 (w/w) MS-grade trypsin/Lys-C protease mix and incubated for 24 h at 37 °C. The reaction was quenched with 1 μL formic acid, peptide digests were enriched using offline reverse-phase liquid chromatography (LC), dried, and reconstituted in a 20 μL solution of 2% acetonitrile and 0.1% formic acid prior to online reverse-phase liquid chromatography tandem mass spectrometry (LC-MS/MS).

**LC-MS/MS analysis:** The LC was carried out on an UltiMate 3000 RSLCnano System (Dionex) fitted with a C18 column (inner diameter = 75 μm, length = 50 cm; PepMap RSLC). The Mobile phase comprised of 0.1% formic acid (solvent A) and 80% acetonitrile/0.08% formic acid (solvent B). Samples were loaded in solvent A and eluted as follows: 10% B for 5 min, gradient to 55% B over 40 min then to 85% B over 5 min, 85% B for 6 min, followed by 14 min re-equilibration with 2% B. The LC system was coupled to a QTOF Impact II mass spectrometer (Bruker) equipped with an Easy Spray ion source and operated in positive ion mode. The spray voltage was set to 1.5 kV, with 3.0 L/min dry gas and 165 ºC dry temperature, and the full scans were acquired in a TOF MS mass analyzer over a *m/z* 200–2200 range at a spectral rate of 2.0 Hz. The auto MS/MS analyses with a fixed precursor cycle time of 3s were performed using collision induced dissociation (CID). The precursor was released after 0.3 min. The raw files were converted to .mgf format by the data analysis software (Bruker Daltonik) and were searched against reported proteomes using the ProteinScape software with an in-house Mascot search engine (Matrix Science Inc., Boston, MA). The search parameters were set as follows: peptide tolerance, 20 ppm; MS/MS tolerance, 0.5 Da; enzyme, trypsin; 2 missed cleavage allowed; and fixed carbamidomethyl modifications of cysteine. Oxidation of methionine and protein N-terminal acetylation were used as variable modifications.

Label-free protein quantification was done using the MaxQuant software (version 1.6.5.0) with default parameters[2,23]. The raw data was searched against the Universal Protein Resource (UniProt) database using the Andromeda search engine[24].

**Cell culture.** Cell lines were purchased from American Type Culture Collection (ATCC). Prior to use, the cells were authenticated and tested for mycoplasma contamination by Charles River Laboratories (Margate, United Kingdom). Murine breast cancer 4T1 cells (ATCC no. CRL-2539) and human monocytic leukemia THP-1 cells (ATCC no. TIB-202) were cultured in RPMI 1640 medium supplemented with 10% FBS, 4 mM L-glutamine, 1 mM sodium pyruvate and 1% penicillin/streptomycin, in 5% $CO_2$ at 37 °C.

**Cancer cell uptake of ALUMSNs.** Intracellular imaging and flow cytometry analysis were used for the pH dependent uptake analysis of Ce6-ALUMSNs as previously reported[25]. Briefly, 4T1 cells were seeded at a density of $2\times10^5$ cells/well in 500 µL complete medium in 4-chambered 35 mm glass bottom Cellview cell culture dishes (Greiner Bio-One, Monroe, NC, USA). After culturing for 24 h, the medium was replaced with fresh medium (pH 6.5 or 7.4) containing Ce6-ALUMSNs (0.5 µg/mL Ce6) and incubated at 37 °C under 5% $CO_2$ for 1–4 h. Then, the media in the chambers was replaced with fresh media. Live images were captured under the Olympus FV1000 confocal laser scanning microscope (Cy 5.5 filter) using a 63× Plan-Apo/1.3 NA oil immersion objective with DIC capability. Image processing was done using the Fiji image processing software[26].

In addition to intracellular imaging, pH dependent uptake was determined using the flow cytometry analysis. 4T1 cells were seeded at a density of $1\times10^6$ cells/well in 6-well plates and allowed to grow for 24 h at 37 °C under 5% $CO_2$. Subsequently, the cells were further incubated with Ce6-ALUMSNs (0.5 µg/mL Ce6) for 1 and 4 h. Then, the cells were then washed three times with ice-cold PBS, trypsinized, harvested by centrifugation (1000×*g*, 5 min), washed again twice with ice cold PBS and analyzed by flow cytometry (FACS Aria III, BD Bioscience), with the Cy 5.5 filter.

For cellular uptake pathway analysis, 4T1 cells were seeded at a density of $1\times10^6$ cells/well in 6-well plates and allowed to grow for 24 h at 37 °C under 5% $CO_2$. After 24 h, 4T1 cells were pre-incubated with Ce6-ALUMSNs (0.5 µg/mL Ce6), for 1 h at 4 °C in pH 6.5 serum-free RPMI

1640 media or pretreated for 1 h at 37 °C with 10 mM sodium azide/6 mM 2-deoxy-D-glucose in serum- and glucose-free RPMI, or with the endocytosis inhibitors[27–30] for 30 min at 37 °C. After the addition of Ce6-ALUMSNs (0.5 µg/mL Ce6), the cells were maintained for 1 h at 4 °C or in the presence of inhibitors at 37 °C (10 µM chlorpromazine; 5 mM methyl-β-cyclodextrin; 5 µM filipin; or 5 µM amiloride). Thereafter, the cells were washed three times with ice-cold PBS, trypsinized, centrifuged and re-suspended in 500 µL ice-cold PBS with 10% FBS, and the fluorescence was measured by flow cytometry. Cells treated with Ce6-ALUMSNs without inhibitors were used as control, and cells treated with vehicle alone served as background. The uptake efficiency was determined from the ratio of fluorescence from cells treated with nanospheres under different inhibition conditions to the control cells. Data collection (10,000 cells/sample, gated on live cells by forward/side scatter) was done on a BD FACSAria III cell sorter (BD Biosciences, San Jose, CA), and analysis was performed using the BD FACSDiva software.

**Cell viability**. Cell viability was measured using two complementary assays as previously published[25]. (a) CellTiter 96 AQueous One Solution (MTS) assay[3,31] and (b) Dead Cell Apoptosis assay[32]. Cells were seeded at a seeding density of $5\times10^3$ cells/well in 100 µL complete medium in standard 96-well plates. After culturing for 24 h, the medium was replaced with fresh medium (pH 6.5) containing Ce6-ALUMSNs (0.05–5 µg/mL Ce6) and incubated for 48 h at 37 °C. At 12 h, the cells were exposed NIR light with varying laser irradiation (980 nm, 0.5–1.5 W/cm$^2$) for 3.0 and 5.0 min. Similarly, the biocompatibility of 5–100 µg/mL UMSNs and LUMSNs at pH 7.4 and of 0.05–5 µg/mL Ce6-ALUMSNs at pH 6.5 and 7.4 in the absence of NIR laser irradiation were analysed for 24 h. Thereafter, the medium was replaced with fresh medium, and 20 µL MTS reagent was added to each well. The MTS reagent was incubated for 4 h at 37 °C, and absorbance of the soluble formazan product ($\lambda = 490$ nm) of MTS reduction was measured on a Synergy H1MF Multi-Mode Microplate-Reader (BioTek, Winooski, VT, USA), with a reference wavelength of 650 nm to subtract background. Wells treated with peptide-free carrier were used as control, and wells with medium alone served as a blank. MTS reduction was determined from the ratio of the absorbance of the treated wells to the control wells.

For the Dead Cell Apoptosis Assay, 4T1 cells were seeded at a density of $1\times10^6$ cells/well in 6-well plates and allowed to grow for 24 h at 37 °C under 5% $CO_2$. After 24 h, 4T1 cells were treated with Ce6-ALUMSNs (0.5 µg/mL Ce6) for 12 h at pH 6.5 with varying NIR light irradiation power densities (0.5–1.5 W/cm$^2$). Subsequently, the cells were washed twice with ice-cold PBS, harvested by trypsinization, centrifuged (1000×*g*, 5 min) and re-suspended in 1× annexin-V-binding buffer (10 mM HEPES, 140 mM NaCl, 2.5 mM $CaCl_2$, pH 7.4) to a density of ~$1\times10^6$ cells per mL. The cells were then stained with 5 µL Alexa 488-conjugated annexin V and 0.1 µg

PI per 100 μL of cell suspension for 15 min at room temperature. Immediately afterwards, fluorescence was measured using flow cytometry, and the fractions of live (annexin V-/PI-), early and late apoptotic (annexin V+/PI- and annexin V+/PI+, respectively), and necrotic (annexin V-/PI+) cells were determined.

**Measurement of mitochondrial membrane potential and ROS production**. 4T1 cells were seeded at a density of $2\times10^5$ cells/well in 500 μL complete medium in 4-chambered 35 mm glass bottom Cellview cell culture dishes. After 24 h, 4T1 cells were incubated with Ce6-ALUMSNs (0.5 μg/mL Ce6) for 3 h at pH 6.5 and exposed to NIR laser irradiation (980 nm, 1.5 W/cm$^2$, 3 min). Then, the cells medium were replaced with fresh media and incubated with 200 nM of the mitochondrial membrane potential probe ($\Delta\Psi_m$) TMRM[33] for 30 min and imaged. Imaging was captured on an Olympus Fluoview FV-1000 confocal laser scanning microscope, using a 63× Plan-Apo/1.3 NA oil immersion objective.

The intracellular ROS production was detected by using a DHE probe. 4T1 cells were seeded at a density of of $2\times10^5$ cells/well in 500 μL complete medium in 4-chambered 35 mm glass bottom Cellview cell culture dishes. After 24 h, cells were incubated with Ce6-ALUMSNs (0.5 μg/mL Ce6) for 3 h, irradiated with NIR laser (980 nm, 1.5 W/cm$^2$, 5 min), then replaced with fresh media containing DHE, and imaged under the Olympus FV1000 inverted confocal scanning microscope.

**Macrophage recognition and immunogenicity of the nanospheres.** Differentiated THP-1 cells were seeded at a density of $1\times10^6$ cells/well in 6-well plates and allowed to grow for 24 h at 37 °C under 5% $CO_2$. After 24 h, cells were incubated with Ce6-UMSNs and -ALUMSNs (0.5 μg/mL Ce6) for 4 h. Then, the plates were washed twice with ice cold PBS, harvested by trypsinization, centrifuged (1000×*g*, 5 min) and re-suspended in PBS. The quantification of Ce6-UMSNs and -ALUMSNs uptake was measured using the flow cytometry.

Cell viability of differentiated THP-1 cells treated with Ce6-UMSNs or Ce6-ALUMSNs for 48 h was assessed using the MTS assay. Cells were seeded at a seeding density of $5\times10^3$ cells/well in 100 μL complete medium in standard 96-well plates. After culturing for 24 h, the medium was replaced with fresh medium containing Ce6-UMSNs or Ce6-ALUMSNs (0.05–2 μg/mL Ce6) and incubated for 48 h at 37 °C. Cell viability was assessed using the MTS assay, with the % viability determined from the ratio of the absorbance of the treated cells to the control cells.

The release of inflammatory cytokines, tumor necrosis factor-alpha (TNF-α) and interlukin-1 beta (IL-1β), by macrophages/monocytes once exposed to Ce6-UMSNs and Ce6-LUMSNs was assessed. Differentiated THP-1 cells were seeded at a density of $2\times10^4$ cells/well in

100 μL complete medium in standard 96-well plates. After culturing for 24 h, the medium was replaced with medium containing Ce6-UMSNs and Ce6-ALUMSNs (0.5 μg/mL Ce6) for 24 h at pH 7.4. Cells treated with the macrophage activator LPS[34] were used as a positive control, while untreated cells served as a negative control. Thereafter, the cell culture medium was assayed for secretion of TNF-α and IL-1β using commercial ELISA kits. The total TNF-α and IL-1β levels were determined from the absorbance ($\lambda = 450$ nm) measured on a BioTek Synergy H1MF Multi-Mode Microplate-Reader using a standard TNF-α concentration calibration curve.

***In vivo* studies.** All animal experiments were approved by the NYU Abu Dhabi Institutional Animal Care and Use Committee (NYUAD-IACUC; Protocol No. 21-0005), and were carried out in accordance with the Guide for Care and Use of Laboratory Animals. Female BalbC nude mice, 8 weeks old, were obtained from Jackson Laboratories in Bar Harbor, ME, USA and bred in-house at the NYU Abu Dhabi Vivarium Facility on a 12 h light/dark schedule.

To conduct pharmacokinetics, thermal imaging, biodistribution studies and tumor inhibition, $2\times10^5$ viable breast cancer 4T1 cells were injected subcutaneously into the right flank of each BalbC nude mice. Mice were assessed daily for overt signs of toxicity. Tumor volume was measured via high-precision calipers (Thermo Fisher) using the following formula:

$$\text{Tumor volume (mm}^3\text{)} = (W^2\times L)/2 \qquad (\textit{Equation 2})$$

where W and L are tumor width and length in mm, respectively. Mice were euthanized once tumor volume approached burden defined by NYUAD-IACUC.

To determine the pharmacokinetics of Ce6-ALUMSNs, both plasma and tumor tissues were used to extract Ce6. The concentration of Ce6 in the plasma of mice was quantified using a high-performance liquid chromatography (HPLC) method[19]. Once the 4T1 tumor volume reached 75 mm$^3$, the mice ($n = 4$ per group) were treated with either Ce6 (2.5 mg/kg) or 11 mg/kg Ce6-ALUMSNs (2.5 mg/kg Ce6) via intravenous (*i.v.*) injection. Blood samples were collected at different time points over 48 h via terminal cardiac puncture using K3-EDTA as an anticoagulant under $CO_2$ anesthesia. The collected blood was then processed for plasma by centrifugation (1500×*g*, 5 min), and 80 μL of plasma was separated from whole blood by centrifugation (1500×*g*, 5 min) and spiked with Ce6 (1 μg/mL). Tumor tissue samples were collected from mice ($n = 4$ per group) after 8 h of treatment with either Ce6 (2.5 mg/kg), Ce6-UMSNs, Ce6-LUMSNs, or Ce6-ALUMSNs (2.5 mg/kg Ce6) via intravenous injection. The tumor tissue was homogenized using the Tris buffer (1 M, pH 8), and Ce6 was extracted thrice by dilution in 3 mL chloroform/methanol (9:1 v/v) and vortexed for 10 min, followed by centrifugation (2500×*g*, 10 min). The organic phase from both plasma and tumor tissues was collected and evaporated to dryness under a $N_2$ stream.

The dry residue was then dissolved in 60 µL methanol, and centrifuged (2500×*g*, 10 min). The supernatant was collected and filtered using a 0.2-µm syringe filter. Finally, approximately 20 µL of the supernatant was injected into the HPLC, and peaks were analyzed for Ce6 concentration against a standard calibration curve.

For *in vivo* thermal imaging, 4T1-tumor bearing mice (75 $mm^3$) ($n$ = 4 per group) were *i.v.* injected with: saline, Ce6-UMSNs, Ce6-LUMSNs or Ce6-ALUMSNs (2.5 mg/kg Ce6). At 6 h post-injection, the tumor sites of the mice was irradiated with NIR laser (980 nm, 1.5 $W/cm^2$, 5 min) and images were captured using the infrared camera to record the temperature of the tumor regions of different groups.

To determine the renal clearance of silica, the 4T1 tumor bearing mice (75 $mm^3$) ($n$ = 4 per group) were *i.v.* injected with 11 mg/kg of ALUMSNs (2.5 mg/kg Ce6). At various timepoints (2−72 h) of post *i.v.* injection, urine and feces samples were collected, and Si concentration was determined by ICP–MS against common standards (ICP–MS, Agilent, US)[35].

For *in viv*o tumor reduction studies, 4T1-tumor bearing mice (75 $mm^3$) were randomized into five treatment groups, ($n$ = 16 per group) which were injected intravenously with saline, 11 mg/kg of UMSNs or 11 mg/kg of Ce6-loaded UMSNs, LUMSNs or ALUMSNs (2.5 mg/kg Ce6). Injections were done every 2 days for a total of 15 doses, with the first day of treatment defined as day 0. Within each treatment group, half of the mice were subjected to NIR laser irradiation (980 nm, 1.5 $W/cm^2$, 5 min) at 6 h post injection. Body weight and tumor volume were recorded every 2 days, and survival ($n$ = 4 per group) was monitored for a total of 90 days.

***Ex vivo* $T_1$ mapping.** *Ex vivo* $T_1$ mapping was performed using excised tumor tissue from 4T1-tumor bearing mice (with a tumor volume of 75 $mm^3$; $n$ = 3 per group), which were intravenously injected with either saline, or 11 mg/kg of UMSNs, LUMSNs, or ALUMSNs. Six hours post-injection, the animals were euthanized, and tumors were immediately dissected and imaged using the same imaging procedure that was employed for imaging the solution samples. $T_1$ maps were obtained and quantified in the tumors and, subsequently, the $1/T_1$ relaxation rate was calculated as the reciprocal of $T_1$.

**Histopathology and immunohistochemistry**. Following animal sacrifice at the end of the xenograft experiment on day 30, appropriate sections of the heart, lungs, liver, spleen, kidney and tumor tissues were dissected and fixed in 10% formalin, followed by paraffin embedding for hematoxylin and eosin (H&E)[36,37]. The paraffin embedded tissues were sectioned using the Leica microtome. Around 4 µm thick sections were mounted on glass slides, stained with H&E and examined by light microscopy.

To visualize phenotypic changes during laser induction, IHC analysis[38] was performed. Tumors collected after the conclusion of the treatment were fixed using the 10% formalin followed by paraffin embedding. In brief, the slides were deparaffinized, incubated in 3% methanol-hydrogen peroxide mixtures, followed by treatment with 10 mM EDTA (pH 8.0) at 95 °C. The slides were subsequently cooled to room temperature and rinsed with phosphate buffered saline solution containing 0.05% Tween-20. This was followed by incubation with individual primary antibodies targeting caspase-3 for 1 h. The slides were subsequently rinsed with PBST and incubated with the appropriate HRP-conjugated secondary antibody at room temperature for 30 min. Finally the slides were counterstained with harris hematoxylin, dehydrated in ethanol, mounted with DPX mountant and analyzed under the microscope.

**Statistical analysis**. To ensure unbiased results, all aspects of the *in vitro* experiments including treatment, data acquisition, and data analysis were conducted by different investigators in a blinded manner. Sample sizes for the *in vivo* studies were determined using power calculation based on the NYU Abu Dhabi Institutional Animal Care and Use Committee (NYUAD-IACUC) Protocol (Protocol No. 18-0001). Confidence intervals in this study were calculated based on the standard deviation from at least three biological replicates ($n \geq 3$). Statistical analysis was performed using Prism 7.0 software (GraphPad Software, Inc.; La Jolla, CA, USA). Statistical significance between two groups was determined using an unpaired t-test, while among three or more groups, two-way analysis of variance (ANOVA) followed by Tukey's *post hoc* test was used. A significance level of $P < 0.05$ was considered statistically significant.
